# Motor cortical circuits are uniquely impacted by different exercise intensities

**DOI:** 10.1101/2025.01.24.634315

**Authors:** Nesrine Harroum, Amanda O’Farrell, Layale Youssef, Louna Bohbot, Hajar Maati, Marie Joubert, Benjamin Pageaux, Jason L. Neva

**Affiliations:** École de kinésiologie et des sciences de l’activité physique (EKSAP), Faculté de médecine, Université de Montréal, Montreal, QC, Canada; Centre de recherche de l’Institut universitaire de gériatrie de Montréal (CRIUGM), Montreal, QC, Canada; Centre interdisciplinaire de recherche sur le cerveau et l’apprentissage (CIRCA), Montreal, QC, Canada

**Keywords:** Acute exercise, exercise intensity, neuroplasticity, primary motor cortex

## Abstract

Acute aerobic exercise (AEX) can enhance motor learning and promote neuroplasticity. However, the effect of AEX intensity on primary motor cortex (M1) excitability has not been systematically examined. Hence, the dose-response relationship between AEX intensity and M1 excitability modulation remains unclear. This study investigated the impact of AEX intensity on distinct M1 circuits using transcranial magnetic stimulation (TMS). Thirty right-handed adults underwent four experimental sessions: rest (control), light (LIIT), moderate (MIIT), and high-intensity interval training (HIIT) AEX. AEX intensity was prescribed with the heart rate reserve (HRR) method, and the interval cycling sessions consisted of alternating between 3 min at the target intensity (LIIT: 35% HRR; MIIT: 55% HRR; HIIT: 80% HRR) and 2 min of active recovery (25% HRR) for 20 min total. We performed TMS measures before (Pre), immediately post (Post_0_), and 20 min post (Post_20_) AEX/rest to assess modulation of corticospinal excitability and GABAergic inhibition as measured by short interval-intracortical inhibition (SICI). This study found that: (1) HIIT and MIIT increased corticospinal excitability, with HIIT eliciting a sustained increase; and (2) all AEX intensities (LIIT, MIIT and HIIT) decreased SICI, with the greatest sustained reduction following MIIT. Also, there was a greater reduction in GABAergic inhibition when measured with anterior-posterior than posterior-anterior TMS current following MIIT. Collectively, our results demonstrate the impact of HIIT and MIIT to enhance corticospinal excitability and reduce GABAergic inhibition in M1. This study provides evidence for a dose-response effect of AEX intensity on the modulation of distinct motor cortical circuits.

**KEY POINTS SUMMARY:**

- Acute aerobic exercise (AEX) is known to modulate primary motor cortex (M1) excitability, but the effect of AEX intensity is unclear.
- This study examined the impact of light-, moderate-, and high-intensity interval training (LIIT, MIIT, HIIT) AEX and rest (non-AEX, control) on distinct M1 cortical circuits using transcranial magnetic stimulation (TMS).
- HIIT induced a sustained increase in M1 output excitability, MIIT induced a transient increase, and LIIT showed no effect.
- All exercise intensities (LIIT, MIIT and HIIT) decreased GABAergic inhibition, as measured by short-interval intracortical inhibition (SICI), with MIIT showing a sustained decrease.
- SICI measured with an anterior-to-posterior TMS current demonstrated greater GABAergic disinhibition compared to posterior-to-anterior TMS current following MIIT.
- This study demonstrates a nuanced dose-response impact of AEX intensity on distinct M1 cortical circuits.

## 3. INTRODUCTION

An acute bout of aerobic exercise can enhance motor learning and promote neuroplasticity (Ferrer-Uris et al., 2017; Mang et al., 2014; McDonnell et al., 2013; Neva et al., 2021; Opie & Semmler, 2019; Roig et al., 2012; Stavrinos & Coxon, 2017; Thomas et al., 2016). Studies using transcranial magnetic stimulation (TMS) have shed light on the impact of acute aerobic exercise on the human brain (Andrews et al., 2020; Lulic et al., 2017; Mooney et al., 2016; Neva et al., 2017; Singh et al., 2014; Smith et al., 2014; Yamazaki et al., 2019). A single session of exercise increases the responsiveness to neuroplasticity-inducing repetitive TMS protocols, indicating a potential preparatory or “priming” effect of acute aerobic exercise on neuroplasticity (Andrews et al., 2020; Mang et al., 2014; Singh et al., 2014).

Further investigations have delved into the neurophysiological mechanisms underlying these effects, employing both single- and paired-pulse TMS techniques to explore acute aerobic exercise-induced changes in primary motor cortex (M1) excitability (El-Sayes et al., 2019; Lulic et al., 2017; McDonnell et al., 2013; Mooney et al., 2016; Singh et al., 2014; Smith et al., 2014; Stavrinos & Coxon, 2017; Turco & Nelson, 2021; Yamazaki et al., 2019). Following an acute bout of lower limb cycling exercise, various inhibitory and facilitatory circuits within the upper-limb representation of M1 exhibit excitability changes. Although there is variability in the effects of acute aerobic exercise on these various inhibitory and facilitatory M1 circuits, there are two TMS-based measures that are emerging as more consistent and robust (Youssef et al., 2024). Gathering data shows that acute aerobic exercise decreases GABA_A_-related inhibition, as reflected by a reduction in short-interval intracortical inhibition (SICI; El-Sayes et al., 2019; Lulic et al., 2017; Singh et al., 2014; Smith et al., 2014; Stavrinos & Coxon, 2017; Yamazaki et al., 2019). In fact, a recent meta-analysis demonstrated that decreased SICI was the most consistent and robust effect following acute aerobic exercise, particularly for moderate- and high-intensity exercise (Youssef et al., 2024). Other studies have shown that M1 output excitability, or corticospinal excitability, can be impacted by acute aerobic exercise (El-Sayes et al., 2019; Lulic et al., 2017; MacDonald et al., 2019; Opie & Semmler, 2019; Ostadan et al., 2016). Importantly, even though most studies observed no modulation of corticospinal excitability following acute aerobic exercise across multiple exercise intensities and types (Andrews et al., 2020; El-Sayes et al., 2020; Mang et al., 2014; McDonnell et al., 2013; Morris et al., 2020; Neva et al., 2017; Singh et al., 2014; Smith et al., 2018; Stavrinos & Coxon, 2017; Yamazaki et al., 2019), a meta-analysis demonstrated that only high-intensity cycling exercise increased corticospinal excitability (Youssef et al., 2024). Thus, it appears that acute aerobic exercise parameters, such as intensity, play an important role in the impact of exercise on M1 excitability modulation (Youssef et al., 2024). However, to date, no single study has systematically examined the impact of acute aerobic exercise intensity on M1 cortical circuit modulation, while controlling for exercise type (e.g., continuous vs intermittent) and duration [e.g., 15-30 min; (Youssef et al., 2024)].

The acute aerobic exercise parameters across studies have been considerably different, which may contribute to the variability of the impact of exercise on M1 excitability reported in the literature. Specifically, previous work used different intensities (e.g., light, moderate or high-intensity), types (e.g., continuous or interval exercise) and durations (e.g., 15 minutes, 30 minutes) of exercise, or different combinations of these parameters in a single study (Youssef et al., 2024). For instance, the majority of studies investigating exercise-induced effects on M1 excitability showed decreased SICI using moderate-intensity continuous exercise (El-Sayes et al., 2019; Lulic et al., 2017; Singh et al., 2014; Smith et al., 2014) or high-intensity interval training exercise (HIIT; Hendy et al., 2022; Kuo et al., 2023; Opie & Semmler, 2019; Stavrinos & Coxon, 2017). Only a few studies did not find an effect of SICI following moderate-intensity continuous cycling exercise (Mooney et al., 2016; Morris et al., 2020) or HIIT (Andrews et al., 2020; Nicolini et al., 2020). Only two studies investigated the impact of light-intensity continuous cycling exercise on SICI, which showed inconsistent results (Opie & Semmler, 2019; Yamazaki et al., 2019). On the other hand, most studies that used moderate intensity continuous exercise did not find an effect on corticospinal excitability (Brown et al., 2020; McDonnell et al., 2013; Morris et al., 2020; Neva et al., 2017; Singh et al., 2014; Singh et al., 2016; Smith et al., 2014; Smith et al., 2018), with few studies showing an increase (El-Sayes et al., 2020; Lulic et al., 2017; MacDonald et al., 2019). Some studies have shown that HIIT can increase corticospinal excitability (Hendy et al., 2022; Nicolini et al., 2020; Opie & Semmler, 2019; Ostadan et al., 2016), whereas others have not shown this effect (Andrews et al., 2020; El-Sayes et al., 2020; Mang et al., 2014; Stavrinos & Coxon, 2017). Finally, only few studies investigated the effect of light-intensity continuous exercise on corticospinal excitability and reported no effect (MacDonald et al., 2019; McDonnell et al., 2013; Yamazaki et al., 2019). Thus, it is probable that exercise intensity plays an important role in the impact of acute aerobic exercise on SICI and corticospinal excitability modulation, along with other exercise parameters (i.e., type and duration). Taken together, the current state of the literature supports the potential of a dose-response relationship between exercise intensity and M1 cortical excitability changes. Yet, to date, there is no single study that systematically investigated the impact of acute aerobic exercise on M1 excitability change across a spectrum of exercise intensities (e.g., light, moderate, high), while controlling for exercise type (e.g., continuous or interval) and duration.

It is possible that variability in exercise-induced effects on M1 excitability is due to the TMS parameters used. All previous studies have explored the impact of acute aerobic exercise on M1 excitability changes using a posterior-to-anterior TMS current, except for one previous study (Neva et al., 2021). In that study, both posterior-to-anterior and anterior-to-posterior TMS currents were used to investigate the unique M1 interneurons (Di Lazzaro et al., 2001; Di Lazzaro et al., 2012; Hamada et al., 2014; Hamada et al., 2013; Hanajima et al., 1998; Paulus et al., 2008; Ziemann & Rothwell, 2000) that may be impacted by acute aerobic exercise (Neva et al., 2021). It was found that moderate-intensity cycling exercise increased corticospinal excitability and decreased SICI as measured with anterior-to-posterior, but not with posterior-to-anterior, TMS current. These results suggested that interneuron excitability sensitive to an anterior-to-posterior TMS current may play an important role underlying the exercise-induced effects on neuroplasticity (Hannah et al., 2018; Mirdamadi et al., 2017; Spampinato, 2020; Spampinato et al., 2020). However, no study has investigated the impact of different acute aerobic exercise intensities on M1 interneuron excitability using these different TMS currents (posterior-to-anterior and anterior-to-posterior).

The overarching aim of the current study was to examine the dose-response relationship between acute aerobic exercise and M1 cortical circuit modulation using a single session of interval cycling exercise across a spectrum of different intensities (i.e., light, moderate, high), while holding exercise type (i.e., interval) and duration (i.e., 20 min) constant. The primary objective of this study was to examine the impact of acute aerobic exercise intensity on M1 excitability modulation on SICI and corticospinal excitability. We hypothesized that HIIT would increase corticospinal excitability and that MIIT and HIIT would decrease SICI. The secondary objective was to examine the impact of acute aerobic exercise intensity on M1 cortical circuit excitability as measured by posterior-to-anterior and anterior-to-posterior TMS currents. We hypothesized that corticospinal excitability measured with anterior-to-posterior current would increase following MIIT and HIIT, whereas posterior-to-anterior current would not show this effect. Finally, we expected to observe decreased SICI measured with anterior-to-posterior current to a greater extent than the posterior-to-anterior current following MIIT and HIIT.

## 4. METHODS

### Participants and ethical approval

Characteristics of participants are presented in Table.1. All data are presented as mean (SD) unless otherwise noted. Thirty healthy right-handed (93 [11]; Edinburgh Handedness Inventory [EHI]; Oldfield, 1971) young adults (27 [5] years, 50% females) took part in the study. A sensitivity analysis performed in G*Power with an alpha risk of 0.05 and a sample size of 30 indicated that we had a 90% chance of observing a medium effect size of f(U) = 0.262 (∼ *η^2^_p_* = 0.064) or higher (Lakens, 2022).

Informed consent was obtained before the administration of any experimental protocol. Participants were screened for any potential contraindications to TMS using standard screening forms. Participants reported no neurological disorders and were otherwise healthy. The Ethics Committee of the Centre de recherche de l’Institut Universitaire de Gériatrie de Montréal (CRIUGM) approved all experimental procedures.

### Experimental design

Each participant completed a preliminary visit and 4 experimental sessions with a minimum of 2 days between each session (Fig. 1). The experimental sessions aimed to assess *i)* the impact of acute aerobic exercise (AEX) intensity on M1 excitability, and *ii)* the distinct effects of AEX intensity on M1 excitability measured by posterior-to-anterior (PA) and anterior-to-posterior (AP) TMS currents. Following 5 min at rest to measure resting heart rate, an incremental exercise test to exhaustion was performed during the preliminary visit to obtain peak power output and peak heart rate. Resting heart rate and peak heart rate were used to prescribe the exercise intensity during the subsequent sessions based on heart rate reserve (see heart rate reserve section for more details), and peak power output was used for further categorization of exercise intensity for the AEX sessions. During the 4 experimental sessions, the following four conditions were performed: 20 minutes of *i)* seated rest, *ii)* light-intensity interval training (LIIT), *iii)* moderate-intensity interval training (MIIT), or *iv)* high-intensity interval training (HIIT) AEX. These conditions were performed in a pseudorandomized order, with each session lasting 2-2.5h. Neurophysiological measurements were conducted before (Pre), immediately after (Post_0_), and 20 minutes after (Post_20_) AEX or rest. The order of testing the neurophysiological measures across participants and sessions was pseudorandomized. To accommodate diurnal fluctuations in corticospinal excitability (Merrell et al., 2024), sessions were scheduled at the same time of day (±3 h) for each participant (Tamm et al., 2009).

**Figure 1:**
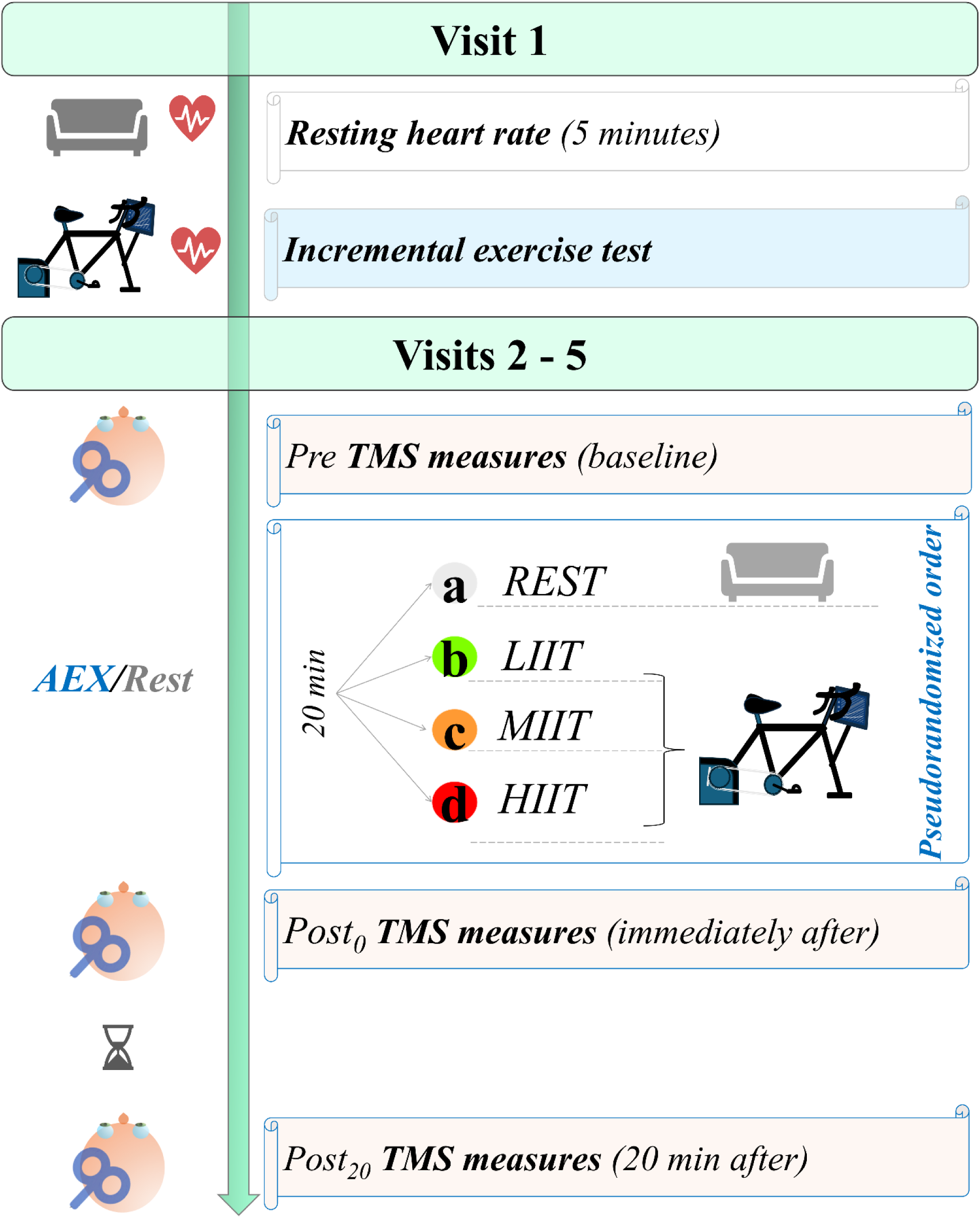
Study Design. An incremental exercise test to exhaustion was conducted in visit 1 (preliminary visit) to establish the exercise intensities for the four subsequent visits (experimental sessions) following ACSM recommendations based on the heart rate reserve. The four experimental sessions involved 20 minutes of either (1) seated rest, (2) light-intensity interval cycling (LIIT), (3) moderate-intensity interval cycling (MIIT), or (4) high-intensity interval cycling (HIIT). The order of each session was randomized. Neurophysiological measurements using transcranial magnetic stimulation (TMS) were taken pre (Pre), immediately post (Post_0_), and 20 minutes post (Post_20_) AEX or rest. <u>Abbreviations</u>: **AEX**: acute aerobic exercise (4 blocs of 3 minutes cycling followed by 2 minutes active recovery at 25% heart rate reserve); **HIIT**: high interval training cycling acute exercise (80% heart rate reserve); **LIIT**: light interval training cycling acute exercise (35% heart rate reserve); **MIIT**: moderate interval training cycling acute exercise (55% heart rate reserve); **Post_0_**: immediately after exercise/rest; **Post_20_**: 20 minutes after exercise/rest; **Pre**: before exercise/rest.

#### International Physical Activity Questionnaire (IPAQ)

Daily levels of physical activity over the 7 days preceding the visit were assessed using the International Physical Activity Questionnaire (IPAQ; Booth, 2000). The IPAQ assesses physical activity across various domains including leisure time, household chores, gardening, as well as work- and transportation-related activities. Frequency and duration of engagement in each of these activities are collected. These activities were classified according to the following categories: *i)* walking, *ii)* moderate-intensity activities and *iii)* high-intensity activities. Physical activity levels among participants were examined to characterize the sample of participants.

#### Incremental test

During the first visit, participants performed an incremental exercise test on an upright cycle ergometer (Cyclus2, CY00100) to establish the AEX intensity for subsequent sessions. The participant started pedaling at a power output equivalent to their body mass (e.g., 70W for 70kg) (ACSM, 2018; Myers et al., 2009). Every 2 minutes, the power output increased by 15, 20, 25 or 30 W, depending on the participant’s body mass. Participants were instructed to maintain a pedaling cadence between 60 and 80 rotations per minute (rpm). They were asked to maintain the same pedaling cadence until exhaustion, defined as being unable to maintain a cadence higher than 60 rpm for 10 s despite verbal encouragement. During the last 20 seconds of each stage, the participant reported their perception of effort intensity using the CR100 scale (Borg & Kaijser, 2006), muscle (thigh) pain using a numerical rating scale ranging from 0 (no pain) to 10 (maximum pain) (Safikhani et al., 2018), and affective response using the Feeling scale (Hardy & Rejeski, 1989).

#### Heart rate (HR) measurement

Prior to and during the incremental test, heart rate (HR) was recorded using a chest belt (Cyclus2) during the exercise and at rest (HR_rest_). We used the percentage of heart rate reserve (%HRR) method to prescribe the cycling exercise intensities. Heart rate reserve (HRR) refers to the difference between the peak heart rate (HR_peak_) recorded during an incremental test and the HR_rest_ measured during 5 minutes of seated rest before the incremental test. To prescribe exercise intensity, we used the Karvonen formula (Garnier et al., 2018; Karvonen & Vuorimaa, 1988):

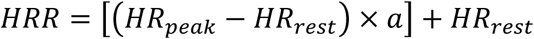

The intensities were prescribed via the constant *a* with the following values corresponding to the exercise intensities as described in the next section: 0.25, 0.35, 0.55, 0.80.

#### Acute Exercise Session

Following a 5-minute warm-up at 25% HRR on the same cycle ergometer used in the incremental exercise test, participants engaged in 20 min of interval cycling, involving four 5-minute blocks. Each block involved 3 min of pedaling at the target intensity of *i)* light (35% HRR, low-intensity interval training [LIIT] condition), *ii)* moderate (55% HRR, moderate-intensity interval training [MIIT] condition), or *iii)* high (80% HRR, high-intensity training [HIIT] condition), followed by 2 min of active rest (25% HRR). These %HRRs were selected according to the American College of Sports Medicine (ACSM) guidelines (ACSM 2018). Participants were instructed to maintain a cycling cadence of 60-80 rpm throughout the acute cycling exercise sessions. Perception of effort using the CR100 scale (Borg & Kaijser, 2006), muscle (thigh) pain perception using a numerical rating scale ranging from 0 to 10 (Safikhani et al., 2018), and affective response using the Feeling scale (Hardy & Rejeski, 1989) were recorded throughout the exercise and rest periods. Specifically, perception of effort, muscle pain, and affective responses were reported by the participants at two time points during each block: *i)* at 0-20 seconds, and *ii)* at 2 minutes 30 seconds, from the beginning of each block. Heart rate was continuously monitored using a heart rate belt (Polar T31C heart rate sensor). At the beginning and end of each cycling exercise interval and the warm-up, the experimenters recorded heart rate. Throughout the cycling exercise, participants were instructed to maintain their hands in a relaxed position, resting them on top of the handlebars and to avoid gripping them. This instruction aimed to minimize contraction of the non-exercised intrinsic hand muscles. Continuous electromyography (EMG) was recorded from both the right and left first dorsal intraosseous (FDI) muscles to ensure participants were resting the non-exercised hand muscles.

#### Rest

The rest condition lasted for 25 minutes to align with the duration of the acute aerobic exercise. Participants sat comfortably on a chair while watching an emotionally neutral documentary. They were instructed not to engage in any tasks involving their upper limbs, such as using a mobile device. Heart rate, quadriceps pain perception using a numerical rating scale ranging from 0 to 10, and affective response using the Feeling scale (Hardy & Rejeski, 1989) were recorded at the same time points as the AEX conditions.

#### Electromyographic recording

For all TMS measurements, EMG was recorded from the right FDI as the muscle of interest and from the right abductor pollicis brevis muscle (APB) to further monitor muscle activity of the hand. Electrodes measuring 1 cm in diameter were placed over the FDI in a belly-tendon configuration. The ground electrode was placed on the ulnar styloid (Covidien, Mansfield, MA, USA). LabChart software (LabChart 8.0) was used to record EMG data. EMG signals were sampled using a PowerLab (PL3516 PowerLab, 16/35 16 Channel Recorder, AD Instruments, Colorado Springs, CO, USA) data acquisition system and a bioamplifier (Dual Bio Amp, AD Instruments, Colorado Springs, CO, USA) with an acquisition rate of 2K Hz and bandpass (20-400 Hz) and notch filtered (center frequency of 50 Hz). Data were captured in a 500-ms sweep from 100 ms before to 400 ms after TMS delivery.

#### Transcranial magnetic stimulation (TMS)

While undergoing TMS measurements, participants were comfortably seated on an adjustable chair and remained at rest. A Magstim BiStim 200^2^ stimulator (Magstim Co., UK) connected to a figure-of-eight coil (Magstim 70 mm P/N 9790, Magstim Co., UK) was used to deliver monophasic TMS pulses. TMS current was manipulated to generate current flow either in a PA or AP direction. A standard coil for PA TMS measures delivered a current that was directed in a posterior to anterior direction. A custom coil was made to produce an AP current, delivering a TMS current in the 180° reverse direction from PA (i.e., anterior-to-posterior). The TMS coils were oriented 45° to the mid-sagittal plane with the handle facing posteriorly. Finally, the standard coil was used to deliver pulses in the lateral-medial (LM) current direction, which was performed by orienting the coil handle 90° to the mid-sagittal plane (Fig. 2; (Cirillo et al., 2018; Di Lazzaro et al., 2001; Hamada et al., 2014; Kaneko et al., 1996; Ni et al., 2011). Coil positioning and monitoring were performed using Brainsight neuronavigation software (Rogue Research Inc., Montreal, QC, Canada). By positioning the coil over M1 in the PA orientation, the “hotspot” for the M1 FDI representation was identified. This hotspot was used for the PA, AP and LM TMS current directions. The resting motor threshold (RMT) was determined at the hotspot for all TMS current directions (PA, AP and LM). RMT was defined as the lowest stimulus intensity needed to elicit 5 out of 10 consecutive MEPs with a peak-to-peak amplitude of 50 μV or more. TMS pulses were administered at a random rate between 0.15 and 0.2 Hz (with ∼20% variation) throughout the study.

**Figure 2:**
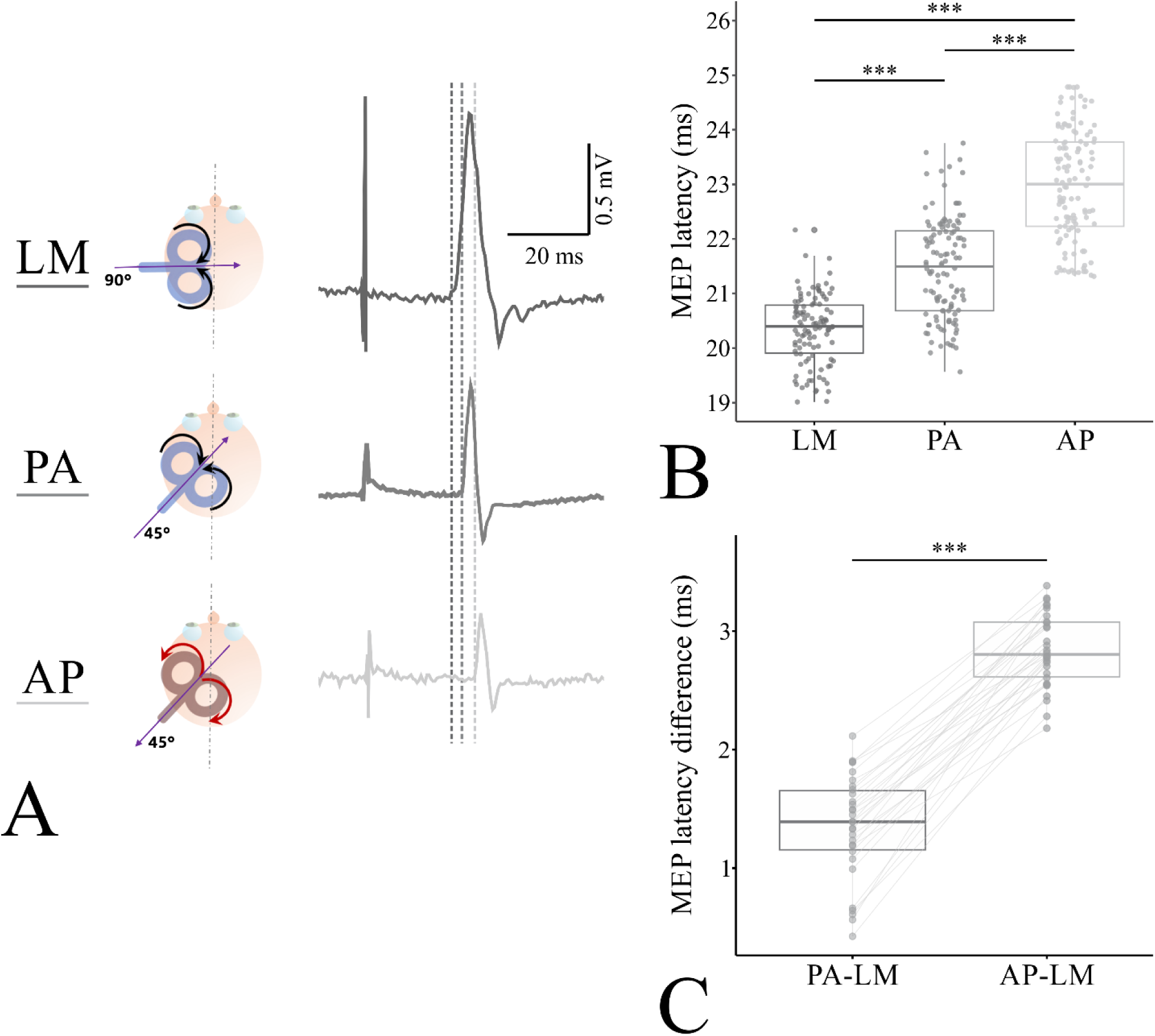
Transcranial magnetic stimulation (TMS) current directions and motor evoked potential (MEP) onset latency results. (A, left panel) TMS current directions are displayed. LM TMS is represented in black, PA TMS is represented in dark grey, and AP TMS is represented in light grey. The figure displays TMS coil current directions, represented with purple arrows, and orientations over the left M1 (dominant) first dorsal intraosseous muscle (FDI) representation. We used a standard 8-figure TMS coil for both LM and PA stimulations, and a reversed current coil for AP stimulations. The coil was oriented 90° for LM TMS current and 45° for both PA and AP TMS currents, relative to the longitudinal fissure. (A, right panel) Electromyographic (EMG) traces are displayed from a representative participant, recorded from the right (dominant) FDI. Vertical dashed green lines represent MEP onset latency elicited by each TMS current direction (LM, PA, AP). (B) MEP onset latency for each TMS current (LM, PA, AP) using boxplots, with dots representing individual data. (C) MEP onset latency differences (LM-PA, LM-AP) using boxplots, with individual data connected with green lines. <u>Abbreviations</u>: AP: anterior-to-posterior; LM: lateral-to-medial; ms: milliseconds; PA: posterior-to-anterior; TMS: transcranial magnetic stimulation; *** p < .001.

MEP amplitudes were evaluated in both the PA and AP current directions to examine potential changes in corticospinal excitability following AEX. MEPs were assessed at 110% and 130% RMT in both PA and AP current directions. MEPs were assessed at 110% RMT since previous work has shown that lower intensity suprathreshold TMS pulses preferentially activate distinct interneuron circuits targeted with PA and AP currents (Cirillo & Byblow, 2016; Cirillo et al., 2018; Day et al., 1989; Di Lazzaro et al., 2001; Hamada et al., 2014; Hamada et al., 2013; Lazzaro et al., 1998; Sakai et al., 1997). Our previous work showed that moderate intensity AEX increased corticospinal excitability measured with this lower intensity (i.e., 110% RMT) in the AP but not the PA TMS current direction, and not at the higher TMS intensities (i.e., 130% RMT and above) in either current direction (Neva et al., 2021). MEPs were also assessed at 130% RMT to reproduce our previous findings that AEX enhances corticospinal excitability assessed in the AP current direction only at the lower stimulus intensities (i.e., 110% RMT) to preferentially activate a distinct group of interneurons (Neva et al., 2021). Here, we aimed to identify distinct intensity-related AEX-induced modulation of these interneuron populations. Ten stimuli were administered at each TMS intensity and current direction, with the order of intensities randomized. The breaks between TMS measurements were about 10-15 seconds, and under 1 minute when switching between PA and AP TMS coils.

SICI was assessed to investigate the effect of AEX intensity on intracortical inhibition, measured in both PA and AP directions, like our previous work (Neva et al., 2021). A subthreshold conditioning stimulus (CS) was delivered prior to a suprathreshold test stimulus (TS) over the M1 FDI hotspot. The CS was delivered at 80% RMT. The TS delivered at a percentage of maximum stimulator output (%MSO) resulting in an average peak-to-peak MEP amplitude of ∼1 mV across 10 stimulations. The interstimulus interval (ISI) between the CS and TS was set at 2 milliseconds (ms) for the PA TMS current direction. The decision to use a 2 ms ISI for PA while measuring SICI was based on previous research indicating a significant reduction in inhibition after acute exercise (Hendy et al., 2022; Lulic et al., 2017; Neva et al., 2021; Opie & Semmler, 2019; Smith et al., 2014; Stavrinos & Coxon, 2017; Yamazaki et al., 2019). Additionally, other studies showed that employing a 3 ms ISI during PA SICI assessment could potentially be affected by facilitation mechanisms (Peurala et al., 2008). Since previous work demonstrated enhanced intracortical facilitatory mechanisms following AEX (Morris et al., 2020; Neva et al., 2017; Smith et al., 2014), this was a particularly important confound to avoid. Conversely, we used a 3 ms ISI to measure SICI in the AP current direction based on previous work that showed consistent and pronounced inhibition compared to a 2 ms ISI (Cirillo et al., 2018; Neva et al., 2021; Sale et al., 2016). Moreover, considering the likely longer cortical transmission pathways with AP TMS (Spampinato, 2020; Spampinato et al., 2020), the 3 ms ISI was deemed appropriate for assessing SICI in this current direction (Neva et al., 2021). To evaluate SICI at each timepoint (Pre, Post_0_, and Post_20_), 10 paired pulses (CS followed by TS) and 10 TS were administered in a randomized order for both PA and AP TMS current directions.

MEP onset latencies were identified as the earliest latency among the block of 10 MEPs to indirectly evaluate I-wave recruitment and infer preferential activation of distinct interneuron populations. This procedure was conducted for both PA and AP TMS current directions (assessed at 110% RMT). Finally, to estimate D-wave activation MEP onset latencies were identified in the same way as described above, expect that it was assessed using an LM TMS current at 150% RMT (Cirillo & Byblow, 2016; Cirillo et al., 2018; Day et al., 1989; Di Lazzaro et al., 2001; Hamada et al., 2014; Hamada et al., 2013; Lazzaro et al., 1998; Sakai et al., 1997).

### Data processing and statistical analysis

Repeated measures analysis of variance (RM-ANOVA) was used to analyze the AEX/rest-related data (HR, perception of effort, muscle pain and affect) and the neurophysiological data as measured using TMS (MEP amplitude, SICI and MEP onset latency). The specific analyses will be described below in more detail. *Post hoc* analyses were conducted using Holm-Bonferroni correction where appropriate. Residual statistics, skewness, and kurtosis values, along with plots, were generated to assess the normality and homoscedasticity of the data. Statistical procedures were performed using Jamovi software (version 2.2.5), with significance set at *p* < .05. Effect sizes were calculated and reported as partial eta squared (*η^2^_p_*) following established guidelines for interpretation (Cohen, 2013), where 0.01 indicates a small effect, 0.06 a moderate effect, and 0.14 a large effect. All data are reported as mean (SD).

#### Aerobic exercise / rest data

We assessed HR, perception of effort, muscle pain and affective response data during the AEX and rest sessions to confirm that the AEX intensities elicited different physiological / psychological responses corresponding to each AEX condition (HIIT, MIIT, LIIT) or rest. Thus, one-way RM-ANOVAs using the within-subjects factor of exercise CONDITION (HIIT, MIIT, LIIT, REST) were conducted separately on average values for HR, perception of effort, muscle pain, and affective response.

#### Neurophysiological data

##### RMT, TS %MSO and TS MEP amplitudes

Consistent RMT (%MSO), determined at the beginning of each session (pre-AEX/rest), was ensured conducting a two-way RM-ANOVA using within-subjects factors CONDITION (HIIT, MIIT, LIIT, REST) and TMS CURRENT (PA, AP). Stable TS %MSO values during SICI assessment pre- and post-AEX/rest were ensured using a three-way RM-ANOVA, considering within-subject factors TIME (Pre, Post_0_, Post_20_), CONDITION (HIIT, MIIT, LIIT, REST) and TMS CURRENT (PA, AP). This ensured stable TMS intensity used during assessment of SICI pre- and post-AEX/rest, across the different AEX conditions and TMS currents. Stable TS MEP amplitudes pre- and post-AEX/rest were ensured using a three-way RM-ANOVA, considering within-subject factors TIME (Pre, Post_0_, Post_20_), CONDITION (HIIT, MIIT, LIIT, REST), and TMS CURRENT (PA, AP). This confirmed consistent corticospinal output excitability throughout the assessment of SICI pre- and post-AEX/rest, across the different AEX conditions and TMS currents.

##### MEP onset latency

We used a semi-automated system to determine MEP onset latency, defined as the point in time where the rectified EMG signal surpassed 5 times the mean pre-stimulus EMG. We calculated MEP latencies induced by single-pulse TMS in each current direction (PA, AP, LM). Additionally, we calculated MEP latency differences (ΔPA-LM, ΔAP-LM) as further indirect indicators of I-wave recruitment, thereby confirming specific interneuron activation as elicited by PA and AP TMS (Cirillo & Byblow, 2016; Cirillo et al., 2018; Di Lazzaro et al., 2001; Hamada et al., 2014; Hamada et al., 2013; Neva et al., 2021; Ni et al., 2011; Sale et al., 2016; Spampinato, 2020). We used the earliest MEP latency response in a two-way RM-ANOVA, incorporating within-subjects factors TMS CURRENT (PA, AP, LM) and CONDITION (HIIT, MIIT, LIIT, REST). Similarly, MEP onset latency differences between PA-LM and AP-LM were compared using a two-way RM-ANOVA, incorporating the within-subjects factor of MEP ONSET DIFFERENCE (ΔPA-LM, ΔAP-LM) and CONDITION (HIIT, MIIT, LIIT, REST).

##### EMG and MEP data processing for corticospinal excitability and SICI

For both MEPs and SICI, we visually inspected the EMG data for voluntary muscle activity. Peak-to-peak MEP amplitudes (measured in millivolts, mV) were analyzed using custom MATLAB scripts. Any trials showing visible voluntary pre-stimulus EMG activity were excluded from the analysis (constituting 0.73% of trials). SICI was quantified as a ratio of CS + TS MEP amplitude over TS MEP amplitude: 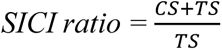, where smaller SICI ratio values indicate greater inhibition, while larger values indicate less inhibition (disinhibition).

##### Corticospinal excitability

To assess the impact of AEX intensity on corticospinal excitability, the average of ten MEP amplitudes was computed for each of the two TMS intensities (110%, 130% RMT). Each stimulus intensity was analysed separately, as the utilization of lower intensities (e.g., 110% RMT) enhances the likelihood of preferentially activating distinct interneuron circuits with both PA and AP TMS current directions (Cirillo & Byblow, 2016; Cirillo et al., 2018; Hamada et al., 2014; Hamada et al., 2013; Mirdamadi et al., 2017; Ni et al., 2011). Thus, we performed three-way RM-ANOVAs using within-subjects factors TIME (Pre, Post_0_, Post_20_), CONDITION (HIIT, MIIT, LIIT, REST), and TMS CURRENT (PA, AP) for each TMS intensity (110, 130% RMT).

##### Short-interval intracortical inhibition

To assess the impact of AEX intensity on SICI, the average MEP amplitude was computed for each of the ten TS and CS+TS pulses, as assessed with each TMS current (PA, AP), to determine the SICI ratio. Thus, a three-way RM-ANOVA was conducted, incorporating within-subjects factors TIME (Pre, Post_0_, Post_20_), CONDITION (HIIT, MIIT, LIIT, REST), and TMS CURRENT (PA, AP) using SICI ratio as the dependent measure.

## 5. RESULTS

### Physical activity data (IPAQ)

Participants demonstrated a large range of physical activity levels as assessed by the long-form IPAQ (Craig et al., 2003). There was a mean (SD) metabolic equivalents-min/week of 4871 (5222) for all participants, with 26.7% (8 participants) categorized as demonstrating high, 26.7% (8 participants) as moderate, and 46.6% (14 participants) as low (see Table.1) levels of physical activity.

### Aerobic exercise/rest data

#### Incremental exercise test

The average incremental test duration was 11.72 (4) minutes. The average peak power output was 173 (44) W. Average peak heart rate was 180 (13) bpm. The average resting heart rate was 67 (10) bpm. The power output used in each exercise condition are illustrated in Fig. 3A as a percentage of peak power output.

**Figure 3:**
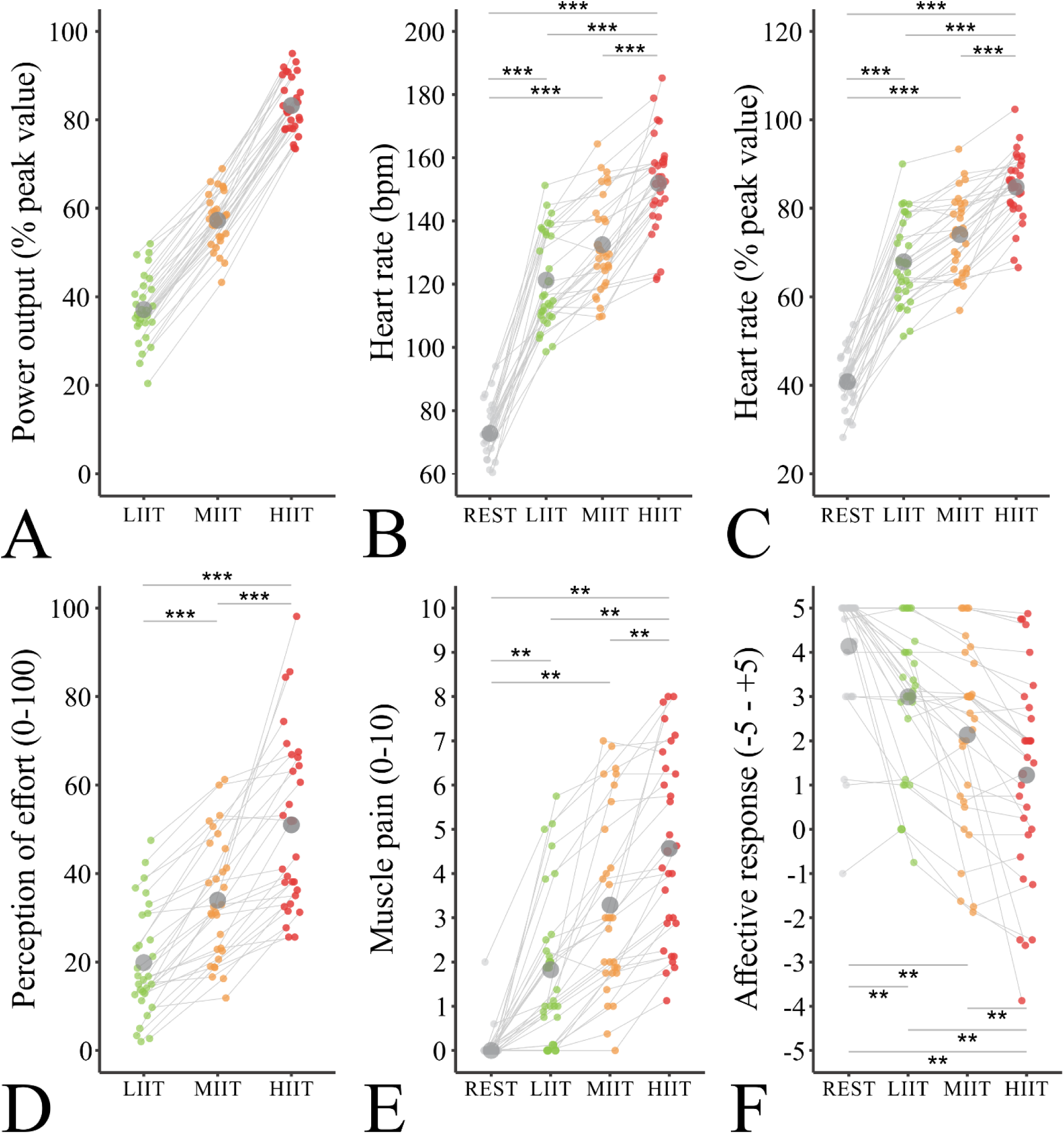
Exercise-related data. (A) Displays the power output normalized to the percent of the peak value reached during the incremental exercise test. (B) Displays the raw heart rate data in bpm. (C) Displays the heart rate data normalized to the percent of the peak value reached during the incremental exercise test. (D) Displays the perception of effort data on CR-100 scale (0-100). (E) Displays the reported pain data on a 0-10 scale. (F) Displays the affective response to exercise assessed using the Feeling Scale. Large dark grey circles represent the average for each condition (REST, LIIT, MIIT and HIIT), and the colored dots represent the individual data. <u>Abbreviations</u>: % peak value: percent of the peak value; bpm: beat per minute; HIIT: high intensity interval training cycling acute exercise; LIIT: light intensity interval training cycling acute exercise; MIIT: moderate intensity interval training cycling acute exercise. ** p < .01; *** p < .001.

#### Acute aerobic exercise/rest sessions

All participants successfully completed the LIIT, MIIT and HIIT sessions (see supporting information – Fig. 1). Based on the incremental test, the light-intensity exercise during LIIT was set at 64 (16) W, the moderate-intensity exercise during MIIT was set at 99 (25) W, and the high-intensity exercise during HIIT was set at 143 (40) W. The active rest intensity set for all exercise conditions (LIIT, MIIT, and HIIT) was set at 46 (14) W. Supporting information – Table 1 displays the HR, perception of effort, and power output outcomes during each exercise/rest condition at different key timepoints. Supporting information – Table 2 and 3 provide the details of the statistical results for the exercise-related data.

**Table 1:** Participant characteristics.

| Characteristic for N = 30 |  |
| --- | --- |
| Sex (Female/Male) | 15/15 |
| Age (years) | 27 (5) |
| Handedness (EHI score) | Right-handed, 93 (11) |
| Weight (kg) | 68 (12) |
| Height (cm) | 171 (9) |
| Body mass index (kg/m<sup>2</sup>) | 23 (3) |
| Peak power output (W) | 173 (44) |
| IPAQ score (total METs) | 4871 (5222) |
| IPAQ categories for levels of physical activity (n/N) |  |
| - High | 8/30 |
| - Moderate | 8/30 |
| - Low | 14/30 |

*All data are reported as Mean (SD) unless otherwise noted. Abbreviations: **BMI**: body mass index; **cm**: centimeters; **EHI**: Edinburgh Handedness Inventory; **IPAQ**: International Physical Activity Questionnaire; **kg**: kilograms; **m<sup>2</sup>**: meters square; **METs**: Metabolic Equivalents; **N**: total number of participants; **n**: number of participants in the indicated sub-group; **W**: Watts.*

#### Raw heart rate data

HR values (see Fig. 3B) showed a significant effect of CONDITION (F_3,87_ = 268.07, *p* < .001, *η^2^_p_* = 0.902), with post hoc analyses demonstrating significant differences between all conditions (all *ps* < .001). Specifically, HIIT showed the highest HR (151 [15] bpm), followed by MIIT (131 [7] bpm), LIIT (122 [4] bpm) and REST (73 [1] bpm).

#### Heart rate data as a percentage of peak value

For HR (% peak value; see Fig. 3C), a main effect of CONDITION (F_3,87_ = 361.5, *p* < .001, *η^2^_p_* = 0.93) was found. Post hoc analysis indicated that all conditions were different from each other (all *p*s < .001). Specifically, HIIT showed the highest peak % HR (85 [8] bpm), followed by MIIT (74 [9] bpm), LIIT (68 [10] bpm) and REST (41 [6] bpm).

#### Perception of effort data

For the perception of effort (Fig. 3D), a main effect of CONDITION (F_3,58_ = 51.56, *p* < .001, *η^2^_p_* = 0.643) was found. Post hoc analysis indicated that all conditions were different from each other (all *ps* < .001). Specifically, HIIT showed the highest perception of effort (51 [14]), followed by MIIT (34 [6]), and LIIT (20 [3]).

#### Muscle pain data

For reported pain (Fig. 3E), a main effect of CONDITION (F_3,87_ = 59.45, *p* < .001, *η^2^_p_* = 0.67) was found. Post hoc analysis indicated that all conditions were different from each other (all *ps* < .002). Specifically, HIIT showed the highest reported pain (4 [1]), followed by MIIT (3 [1]), LIIT (2 [0.4]) and REST (0 [0.02]).

#### Affective response data

For affective responses (Fig. 3F), a main effect of CONDITION (F_3,87_ = 31.54, *p* < .001, *η^2^_p_* = 0.52) was found. Post hoc analysis indicated that all conditions were different from each other (all *ps* < .001). Specifically, HIIT showed the lowest score on the Feeling Scale (1 [1]), followed by MIIT (2 [0.4]), LIIT (3 [0.2]) and REST (4 [0.1]).

### Neurophysiological data

#### Baseline neurophysiological data

For RMT values, a main effect of CURRENT (F_2,58_ = 81.602, *p* < .001, *η^2^_p_* = 0.74) was found (see supporting information – Table 4). Post hoc analysis revealed that RMT for PA TMS current (45 [6] %MSO) was significantly lower than both AP (57 [9] %MSO; *t_29_* = −5.743, *p* < .001) and LM (51 [60] %MSO; *t_29_* = −7.173, *p* < .001) TMS currents. Further, RMT values for LM current was significantly lower than AP TMS current (*t_29_* = −5.743, *p* < .001). There was no CONDITION main effect (F_3,87_ = 0.665, *p* = .576, *η^2^_p_* = 0.02) or CURRENT × CONDITION interaction (F_6,174_ = 0.408, *p* = .873, *η^2^_p_* = 0.01).

TS %MSO values used during SICI were stable across time and exercise conditions but were different between TMS currents (see supporting information – Table 5). Specifically, there was a main effect of CURRENT (F_1,294_ = 402.61, *p* < .001, *η^2^_p_* = 0.93), no significant effect of TIME (F_2,58_ = 0.510, *p* = .650, *η^2^_p_* = 0.02), CONDITION (F_3,87_ = 0.3, *p* = .822, *η^2^_p_* = 0.01), TIME × CONDITION interaction (F_6,174_ = 1.35, *p* = .239, *η^2^_p_* = 0.04) or TIME × CONDITION × CURRENT interaction (F_6, 174_ = 1.07, *p* = .379, *η^2^_p_* = 0.04).

TS MEP amplitudes during SICI were stable across time and exercise conditions but were different between TMS currents (see supporting information – Table 6). Specifically, there was a main effect of CURRENT (F_1,294_ = 402.61, *p* = .001, *η^2^_p_* = 0.93), no significant effect of TIME (F_2,58_ = 0.23, *p* = .797, *η^2^_p_* = 0.01), CONDITION (F_3,87_ = 2.33, *p* = .079, *η^2^_p_* = 0.07), nor was there CONDITION × CURRENT interaction (F_3,87_ = 0.68, *p* = .567, *η^2^_p_* = 0.02) or TIME × CONDITION × CURRENT interaction (F_6,174_ = 1.35, *p* = .239, *η^2^_p_* = 0.04).

For MEP onset latency (see Fig. 2), a significant main effect of CURRENT (F_2,58_ = 243.81 *p* < .001, *η^2^_p_* = 0.91) was found. Post-hoc analysis showed that MEP onset latency was shorter using LM current (20 [1] ms) compared to PA (21 [1] ms; *t_29_* = −11.06, *p* < .001) and AP TMS current (23 [1] ms; *t_29_* = −24.29, *p* < .001). Further, MEP onset latency for using PA TMS current was shorter than AP TMS current (*t_29_* = −14.47, *p* < .001). There was no significant effect of CONDITION or CURRENT × CONDITION interaction (all *ps >* .147).

For MEP onset latency difference (ΔPA-LM, ΔAP-LM; see Fig. 2), statistical analysis revealed a significant effect of MEP ONSET DIFFERENCE (F_1,29_ = 257.51, *p* < .001, *η^2^_p_* = 0.90). Post-hoc analysis showed that ΔPA-LM (2 [1] ms) was greater than ΔAP-LM (3 [1] ms; *t_29_* = −19.15, *p* < .001). There was no significant effect of CONDITION or MEP ONSET DIFFERENCE × CONDITION interaction (all *ps >* .142).

#### Corticospinal excitability

MEP data is displayed for each TMS intensity (Fig. 4A, C and D: 110% RMT; Fig. 4B, E and F: 130% RMT), and TMS currents collapsed (Figs. 4A and B) and separated into PA (Fig. 4C and E: *left panel* average data, *right panel* individual data plots with means overlayed) and AP (Fig. 4D and F: *left panel* average data, *right panel* individual data plots with means overlayed). For MEPs at 110% RMT (Fig. 4.A), a TIME × CONDITION interaction was found (F_1,29_ = 22.16, *p* < .001, *η^2^_p_* = 0.2). Post hoc analysis indicated greater MEP amplitude at Post_0_ compared to Pre for HIIT (*t_29_* = −6.42, *p* < .001) and MIIT (*t_29_* = −4.41, *p* = .008). At Post_0_, MEP amplitude was greater for HIIT (*t_29_* = 4.88, *p* < .001) and MIIT (*t_29_* = 3.42, *p* = .047) compared to rest. Also, there was greater MEP amplitude at Post_20_ compared to Pre for HIIT (*t_29_* = −5.5, *p* < .001). Finally, at T2, MEP amplitude was greater for HIIT compared to rest (*t_29_* = 4.06, *p* = .009). Additionally, we found main effects of TIME (F_2,58_ = 27.64, *p* < .001 *η^2^_p_* = 0.49), CONDITION (F_3,87_ = 7.53, *p* < .001, *η^2^_p_* = .21), CURRENT (F_3,87_ = 7.53, *p* < .001, *η^2^_p_* = 0.43), with no other effects or interactions (all *ps* < .106).

**Figure 4:**
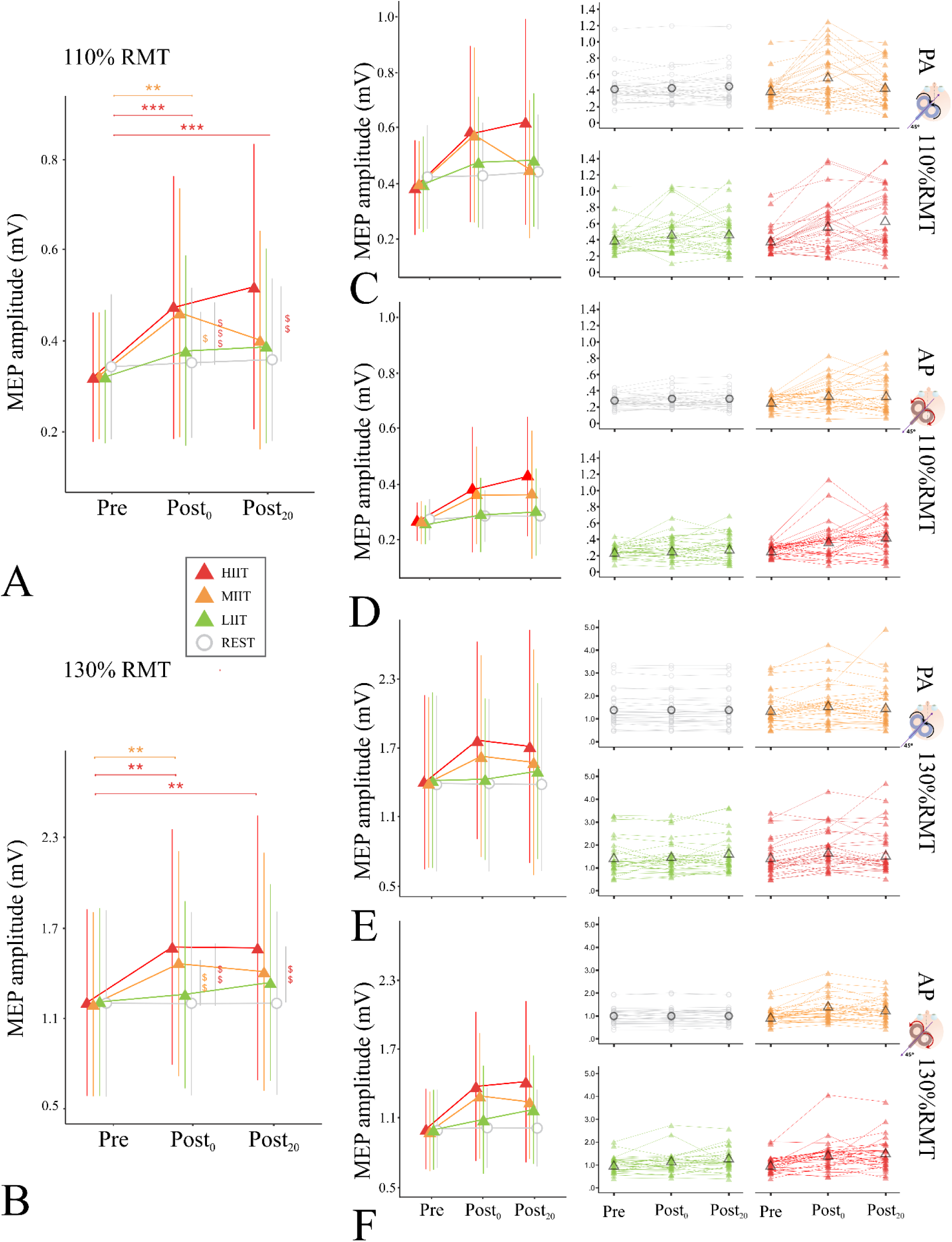
Corticospinal excitability results. **(A, B)** Display average peak-to-peak MEP amplitudes elicited with TMS currents collapsed for 110% RMT **(A)** and 130% RMT **(B)** at each timepoint (Pre, before exercise/rest; Post_0_, immediately after exercise/rest; Post_20_, 20 minutes after exercise/rest). Vertical grey lines show significant differences between rest and the condition represented by its color, with green for LIIT, orange for MIIT, and red for HIIT average peak-to-peak MEP amplitudes elicited with PA current for 110% RMT **(C, left panel)**, AP current for 110% RMT **(D, left panel)**, PA current for 130% RMT **(E, left panel)** and AP current for 130% RMT **(F, left panel)** are displayed at each timepoint. **(Right panels of C, D, E, F)** Box plots display individual data at each timepoint for MEP amplitudes at 110% RMT elicited with PA and AP currents, and for 130% RMT with PA and AP currents, in which the box depicts the median, 25th and 75th percentiles, and the individual data is overlayed. HIIT is displayed in red triangles, MIIT is displayed in orange triangles, LIIT is displayed in green triangles, and Rest is displayed in gray rings. Bars represent the standard deviation of the mean. <u>Abbreviations:</u> **AP**: anterior-to-posterior; **HIIT**: high intensity interval training cycling acute exercise; **LIIT**: light intensity interval training cycling acute exercise; **MEP**: motor evoked potential; **MIIT**: moderate intensity interval training cycling acute exercise; **mV**: millivolt; **RMT**: resting motor threshold; **PA**: posterior-to-anterior; **Post_0_**: immediately after exercise/rest; **Post_20_**: 20 minutes after exercise/rest; **Pre**: before exercise/rest; **TMS**: transcranial magnetic stimulation. Within-subjects: * p < .05; ** p < .01; **\*\*\*** p < .001. Between-subjects: $ p < .05; $$ p < .01; **$$$** p < .001.

For MEPs at 130% RMT (Fig. 4.B), a TIME × CONDITION interaction was found (F_1,29_ = 8.43, *p* < .001, *η^2^_p_* = 0.23). Post hoc analysis indicated greater MEP amplitude at Post_0_ compared to Pre for HIIT (*t_29_* = −5.97, *p* = .001) and MIIT (*t_29_* = −5.32, *p* = .007). At Post_0_, MEP amplitude was greater for HIIT (*t_29_* = 6.05, *p* = .001) and MIIT (*t_29_* = 4.84, *p* = .001) compared to rest. Also, there was greater MEP amplitude at Post_20_ compared to Pre for HIIT (*t_29_* = −5.82, *p* = .002). Finally, at Post_20_, MEP amplitude was greater for HIIT compared to rest (*t_29_* = 5.98, *p* = .001). Additionally, we found main effects of TIME (F_2,58_ = 25.27, *p* < .001 *η^2^_p_* = 0.47), CONDITION (F_3,87_ = 17.98, *p* < .001, *η^2^_p_* = 0.38), CURRENT (F_3,87_ = 8.4, *p* = .007, *η^2^_p_* = 0.22), with no other effects or interactions (all *ps* < .577).

#### Short-interval intracortical inhibition

SICI data is displayed for TMS currents collapsed (Fig. 5A) and separated into PA (Fig. 4B: *left panel* average data, *right panel* individual data plots with means overlayed) and AP (Fig. 4D: *left panel* average data, *right panel* individual data plots with means overlayed). For SICI (Fig. 5A), a TIME × CONDITION interaction was found (F_6,174_ = 7.28, *p* < .001, *η^2^_p_* = 0.2). Post hoc analysis indicated less SICI at Post_0_ compared to Pre for HIIT (*t_29_* = −7.38, *p* < .001), MIIT (*t_29_* = −6.06, *p* < .001) and LIIT (*t_29_* = −4.88, *p* < .001). Also, there was less SICI at Post_20_ compared to Pre for HIIT (*t_29_* = −5.38, *p* < .001), MIIT (*t_29_* = −5.77, *p* < .001) and LIIT (*t_29_* = −3.68, *p* = .022). At Post_0_, there was less SICI for HIIT (*t_29_* = 4.58, *p* =.002) and MIIT (*t_29_* = 3.67, *p* = .023) compared to rest. Finally, at Post_20_, there was less SICI for MIIT compared to rest (*t_29_* = 3.6, *p* = .026).

**Figure 5:**
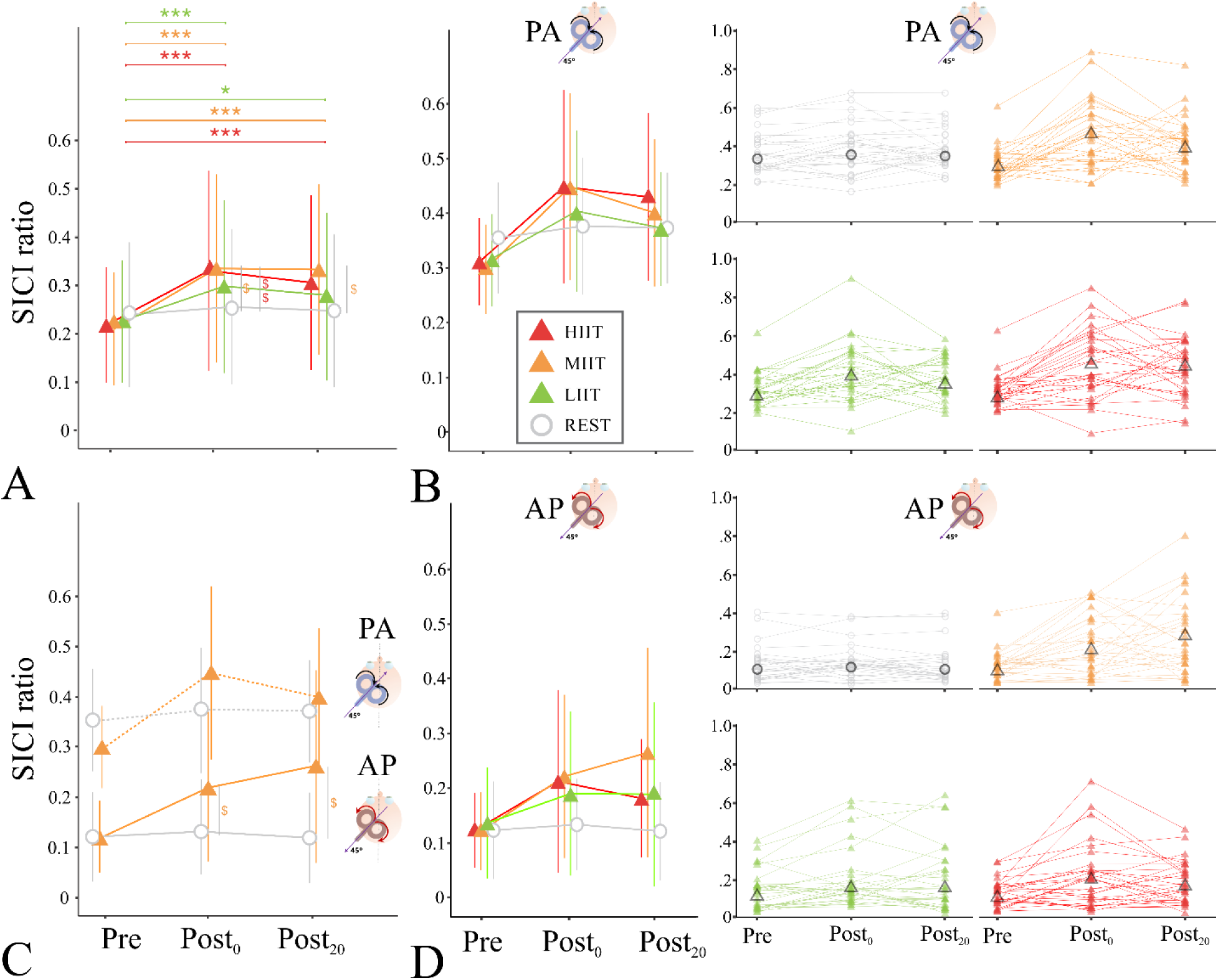
Short-interval intracortical inhibition results. **(A)** Displays average SICI ratios, where greater values represent less GABAergic inhibition, at each timepoint (Pre, before exercise/rest; Post_0_, immediately after exercise/rest; Post_20_, 20 minutes after exercise/rest). Vertical grey lines show significant differences between rest and the condition represented by its color, with green for LIIT, orange for MIIT, and red for HIIT. **(Left panels of B, D)** average SICI ratios at each timepoint elicited with PA **(B, left panel)** and AP **(D, left panel)** currents. **(Right panels of B, D)** Box plots for SICI ratios display individual data at each timepoint elicited with PA **(B, right panel)** and AP **(D, right panel)** currents, in which the box depicts the median, 25th and 75th percentiles, and the individual data is overlayed. HIIT is displayed in red triangles, MIIT is displayed in orange triangles, LIIT is displayed in green triangles, and Rest is displayed in gray rings. Bars represent the standard deviation of the mean. <u>Abbreviations</u>: AP: anterior-to-posterior; HIIT: high intensity interval training cycling acute exercise; LIIT: light intensity interval training cycling acute exercise; MIIT: moderate intensity interval training cycling acute exercise; PA: posterior-to-anterior; Post_0_: immediately after exercise/rest; Post_20_: 20 minutes after exercise/rest; Pre: before exercise/rest; SICI: short intracortical inhibition; TMS: transcranial magnetic stimulation. Within-subjects: * p < .05; ** p < .01; **\*\*\*** p < .001. Between-subjects: $ p < .05; $$ p < .01; **$$$** p < .001.

We also found a CONDITION × CURRENT interaction (F_3,87_ = 2.83, *p* = .043, *η^2^_p_* = 0.09; Fig. 5C). Post hoc analyses demonstrated that MIIT showed less SICI compared to rest with AP current (*t_29_* = 4.07, *p* = .005), but not PA TMS current (*t_29_* = 0.8, *p* = 0.429). We conducted follow-up pairwise comparisons to understand what was driving this particular effect (Fig. 5B; see supporting information – Table 7). We found that MIIT showed less SICI than rest at Post_0_ and Post_20_ using AP TMS current (all *ps* < .0168), but not with PA current (all *ps* > .490), and there were no differences at Pre for either TMS current (all *ps* > .057; Holm-Bonferroni correction applied for 16 comparisons). Finally, we found a main effect of TIME (F_2,58_ = 48.71, *p* = .001, *η^2^_p_* = 0.63), CONDITION (F_3,87_ = 5.73, *p* < .001, *η^2^_p_* = 0.16), and CURRENT (F_1,29_ = 106.42, *p* < .001, *η^2^_p_* = 0.79).

## DISCUSSION

We investigated the impact of acute aerobic exercise intensity on distinct motor cortical circuits. There are two main findings of the current study: (1) HIIT and MIIT increased corticospinal excitability, with HIIT eliciting a sustained increase in the excitable output of M1; and (2) LIIT, MIIT and HIIT demonstrated a reduction in GABAergic inhibition as measured by SICI, with a sustained reduction after MIIT. A secondary finding was that GABAergic inhibition, as measured by SICI, showed a greater reduction when assessed with anterior-to-posterior than posterior-to-anterior TMS current following MIIT compared to rest. Collectively, the current results demonstrate the capacity of an acute session of high- and moderate-intensity interval exercise to enhance corticospinal excitability and reduce GABAergic inhibition of the motor cortex. These results provide evidence for a dose-response effect of exercise intensity on the modulation of distinct motor cortical circuits.

We collected physiological and perceptual responses during the acute aerobic exercise. Heart rate, perception of effort and muscle pain were higher during the three exercise conditions compared to rest, and affective responses were lower during the three exercise conditions compared to rest. Second, there were clear distinct physiological and perceptual responses between the three exercise conditions. We observed an increased heart rate, perception of effort and muscle pain with increased exercise intensity. Additionally, affective responses decreased with increasing exercise intensity. These differences between conditions across our variables attest that we were successful in our experimental manipulation of three distinct exercise intensities.

### Increased corticospinal excitability following acute exercise

We found that HIIT increased corticospinal output excitability to the greatest extent and duration relative to the other exercise intensities (MIIT and LIIT) and rest. Both MIIT and HIIT increased corticospinal excitability immediately after exercise, but only HIIT maintained this increase for 20 minutes post-exercise. Further, HIIT alone showed increased corticospinal excitability when compared to rest 20 minutes post-exercise. Importantly, these results were found regardless of the measured TMS intensity (110%, 130% RMT) and current direction (posterior-to-anterior, anterior-to-posterior). These results suggest that intensity plays an important role in the ability of acute exercise to modulate the excitability of motor cortex output. In line with the current results, our recent systematic review and meta-analysis demonstrated that only high-intensity exercise increases corticospinal excitability, whereas moderate- and light-intensity exercise does not (Youssef et al., 2024). However, a limitation of this meta-analysis was that each study performed different acute exercise types, durations and intensities (e.g., % maximum power output, max heart rate). Relatedly, no study systematically examined the impact of acute exercise on M1 excitability across a spectrum of exercise intensities (e.g., light, moderate, high), while also simultaneously controlling for exercise type (e.g., continuous vs. interval) and duration. Importantly, we addressed these limitations by examining the effect of acute exercise across three different intensities (i.e., light, moderate, high; based on ACSM guidelines) along with a rest condition, while controlling for exercise type (i.e., interval training exercise) and duration (i.e., 20 min). Thus, the present study represents an experimental confirmation of several results in our meta-analysis (Youssef et al., 2024).

In fact, the current results further extend our previous findings (Youssef et al., 2024) by demonstrating that HIIT increased corticospinal excitability for up to 20 min following exercise compared to the other exercise conditions (i.e., MIIT and LIIT). Evidence has been accumulating to demonstrate that HIIT can increase corticospinal excitability of the non-exercised upper limb (Hendy et al., 2022; Nicolini et al., 2020; Ostadan et al., 2016). The current findings suggest that acute exercise intensity, along with other exercise parameters such as type and duration, are important factors influencing the impact of acute exercise on M1 excitable output. Previous work showed increased corticospinal excitability following HIIT when the intensity was high compared to similar studies (e.g., 105-120% VO_2peak_) and exercise duration was relatively short (e.g., 15 min) (Hendy et al., 2022; Nicolini et al., 2020; Ostadan et al., 2016). In contrast, other studies with slightly lower intensities than the previous examples (Hendy et al., 2022; Nicolini et al., 2020; Ostadan et al., 2016) did not observe an effect of exercise on corticospinal excitability (Andrews et al., 2020; El-Sayes et al., 2020; Mang et al., 2014; Stavrinos & Coxon, 2017). Overall, this suggests that exercise parameters like exercise intensity, type and duration play a key role in exercise-enhanced corticospinal excitability.

Contrary to our hypothesis, MIIT increased corticospinal excitability immediately post-exercise. While our meta-analysis showed that moderate-intensity exercise did not change corticospinal excitability (Youssef et al., 2024), there are individual studies that suggested moderate-intensity exercise can increase corticospinal excitability (El-Sayes et al., 2019; MacDonald et al., 2019). However, it is important to highlight that the majority of studies using moderate-intensity exercise did not find an effect on corticospinal excitability (Andrews et al., 2020; Brown et al., 2020; El-Sayes et al., 2020; Kuo et al., 2023; McDonnell et al., 2013; Morris et al., 2020; Neva et al., 2017; Neva et al., 2021; Singh et al., 2014; Smith et al., 2014; Smith et al., 2018). It may be that our current findings relate to employing MIIT (i.e., moderate-intensity *interval* training exercise) as opposed to all but one study using moderate-intensity *continuous* exercise (El-Sayes et al., 2020). Notably, the vast majority of studies using moderate continuous cycling exercise did not show an impact of acute aerobic exercise on corticospinal excitability (Brown et al., 2020; Kuo et al., 2023; McDonnell et al., 2013; Morris et al., 2020; Neva et al., 2017; Singh et al., 2014; Singh et al., 2016; Smith et al., 2014). The moderate intensity exercise parameters used in the current study, such as incorporating active recovery intervals instead of cycling at a constant load, might partially explain the differing findings with previous work using moderate-intensity continuous exercise (Brown et al., 2020; Kuo et al., 2023; McDonnell et al., 2013; Morris et al., 2020; Neva et al., 2017; Singh et al., 2014; Singh et al., 2016; Smith et al., 2014). In fact, the only study that used MIIT showed increased corticospinal excitability following exercise (El-Sayes et al., 2020). Studies comparing continuous versus intermittent cycling at a moderate intensity have shown that MIIT results in higher blood lactate levels compared to continuous exercise (Beneke et al., 2003; Grossl et al., 2012; Kanthack et al., 2020). It was shown that performing moderate aerobic exercise in intervals (i.e., alternating lower and higher exercise intensities) leads to a lack of stabilization of physiological changes for the same duration as continuous exercise, resulting in greater blood lactate accumulation (Beneke et al., 2003; Grossl et al., 2012; Kanthack et al., 2020). This has been suggested to trigger signaling cascades that increase M1 excitability (Coco et al., 2014; Coco et al., 2010; Coco et al., 2020). Since the current study did not measure these specific physiological changes (e.g., blood lactate), the mechanisms underlying these exercise-induced changes in corticospinal excitability remain speculative.

### Short-interval intracortical inhibition decreased following acute exercise

Our findings indicate a dose-response relationship between exercise intensity and modulation of SICI. All three exercise conditions (HIIT, MIIT and LIIT) decrease SICI immediately and 20 min following exercise compared to pre-exercise. However, only HIIT and MIIT decreased SICI to a greater extent than rest immediately following exercise, and only MIIT showed a decrease in SICI compared to rest 20 min post-exercise. Thus, while each of our exercise intensities were sufficient to modulate SICI, there was a nuanced effect that includes a greater decrease immediately following exercise for HIIT and MIIT than LIIT, with MIIT demonstrating the greatest decrease in SICI. Exercise-induced decreases in SICI have been observed in many studies (El-Sayes et al., 2019; Hendy et al., 2022; Kuo et al., 2023; Lulic et al., 2017; Neva et al., 2021; Opie & Semmler, 2019; Singh et al., 2014; Smith et al., 2014; Stavrinos & Coxon, 2017), and was revealed to be the most consistent and robust effect in our meta-analysis (Youssef et al., 2024). Our moderator analysis showed that moderate- and high-intensity acute exercise contributed to this decrease in SICI, which lasted up to 30 min post-acute aerobic exercise (Youssef et al., 2024). The current results confirm and extend these findings, by showing that MIIT and HIIT reduce SICI immediately following exercise, with MIIT demonstrating a prolonged effect up to 20 min post-exercise compared to rest. Our findings also extend our previous findings by demonstrating that LIIT was also sufficient to elicit a decrease in SICI up to 20 min post-exercise. While our meta-analysis demonstrated that light-intensity exercise did not show decreased SICI, it is likely that the low number of studies employing this intensity contributed to the lack of effect (Youssef et al., 2024). Studies demonstrating exercise-induced disinhibition of SICI hypothesized that this effect is associated with modulation of GABA_A_-receptor mediated activity. Previous pharmacological studies have demonstrated the role of these receptors in the paradigm of SICI measured using TMS (Di Lazzaro et al., 2007; Di Lazzaro et al., 2012; Hanajima et al., 1998). Therefore, exercise-induced changes in SICI might result from a series of physiological events from acute aerobic exercise that lead to modulation of GABAergic functioning within M1.

Both blood lactate and dopamine levels may play a role in the exercise-induced reduction of GABAergic activity underlying SICI. Specifically, blood lactate accumulation is suggested to be associated with a reduction in GABAergic neuronal activity (Kann, 2024), which might partly explain the reduction in SICI following HIIT and MIIT. Similarly, higher exercise intensities (moderate, high) are shown to induce greater levels of dopamine in the brain (Greenwood, 2019; Sacheli et al., 2019), and increased dopamine release can contribute to reduced GABA_A_ receptor activity (Flores-Hernandez et al., 2000). Interestingly, recent research has more directly demonstrated the contribution of dopamine receptor activity in exercise-induced modulation of M1 inhibitory circuitry, such as SICI (Curtin et al., 2023). Therefore, the modulation of dopaminergic pathways following acute exercise may contribute to the reduction in SICI observed with HIIT, and particularly the sustained reduction following MIIT, as observed in the current study. Decreased SICI following LIIT suggests that the physiological mechanisms activated by light-intensity exercise are effective at modulating GABAergic functioning in M1, but may not be as potent as those triggered by higher intensities (MIIT and HIIT). Future work is needed to directly investigate the effect of acute aerobic exercise intensity on these underlying neural mechanisms and their contribution to exercise-induced decrease in SICI.

### Effect of acute exercise intensity on specific M1 interneurons

We found a dose-response effect of exercise on SICI as measured by different TMS currents (anterior-to-posterior versus posterior-to-anterior). Specifically, we found a greater decrease in SICI measured with anterior-to-posterior TMS current 20 minutes post-MIIT compared to rest, which was not observed with posterior-to-anterior TMS current. This suggests that our MIIT exercise uniquely impacted M1 interneurons sensitive to anterior-to-posterior TMS current while measuring SICI, like previous findings (Neva et al., 2021). Interestingly, this result has been demonstrated in the current study while using interval exercise at a moderate intensity, while not showing the same effects for HIIT and LIIT. This suggests a potential unique association between M1 interneurons sensitive to anterior-to-posterior TMS current interacting with GABA_A_-receptor related activity that is impacted by MIIT. Interestingly, these same M1 interneurons sensitive to anterior-to-posterior TMS current have been impacted by other interventions as demonstrated using paired-pulse paradigms like SICI (Mirdamadi et al., 2017; Spampinato, 2020). It is also possible that use of anterior-to-posterior current while measuring SICI may be more reliable than posterior-to-anterior current, and thus may be more sensitive to changes to due interventions or specific behaviors (Amassian et al., 1987; Cirillo & Byblow, 2016; Cirillo et al., 2018; Di Lazzaro et al., 2001; Hamada et al., 2014; Lazzaro et al., 1998; Sale et al., 2016). This suggests that SICI measured with anterior-to-posterior TMS current may reveal the acute aerobic exercise effects more sensitively and reliably compared to SICI with posterior-to-anterior TMS current. However, this finding should be interpreted with caution since it was revealed following exploratory analysis.

Contrary to our hypothesis, corticospinal excitability increased following HIIT and MIIT regardless of TMS current direction. We previously found that an acute session of moderate-intensity continuous exercise increased corticospinal excitability as measured with anterior-to-posterior TMS current, but not with posterior-to-anterior TMS current (Neva et al., 2021). Therefore, we did not confirm our previous results (Neva et al., 2021). It is possible that the exercise type played an important role in these effects, as the current study employed interval training exercise (i.e., MIIT), whereas our previous study used continuous cycling exercise at a moderate intensity. The alternating periods of moderate intensity to lower intensity periods of active recovery, may have led to a net positive neural environment that shifted the release of neurotrophic factors and neurotransmitters (Mang et al., 2013; Sale et al., 2008; Skriver et al., 2014) in the motor cortex to increase the excitable corticospinal output of interneurons sensitive to both posterior-to-anterior and anterior-to-posterior TMS currents. However, this remains highly speculative since we did not measure any specific neurotrophic factors and neurotransmitters.

## Limitations

This study has limitations that should be considered when interpreting the results. First, while our primary focus was on the effect of acute aerobic exercise intensity on M1 excitability, we did not control for the total work across different exercise conditions. It is entirely possible that the total work completed during acute aerobic exercise may impact brain excitability. Instead, we controlled for exercise duration to ensure a consistent time frame for comparing different exercise intensity levels (light, moderate, high). Controlling acute aerobic exercise duration attempts to control for differences in outcomes to be related to intensity rather than accumulated fatigue or total energy expenditure (Tschakert & Hofmann, 2013; Zuniga et al., 2011). Second, we chose interval exercise over continuous exercise to ensure that participants could complete all acute aerobic exercise conditions. Particularly during high-intensity cycling, maintaining continuous pedaling for 20 minutes at such intensity could be highly challenging. Yet, it is important to acknowledge that continuous exercise may uniquely influence TMS-based neurophysiological measures across the different acute aerobic exercise intensities. Future studies should investigate the impact of exercise type (interval vs. continuous) on M1 excitability to comprehensively understand the effects of acute aerobic exercise intensity. Third, we used HRR to prescribe exercise intensity. We are aware that this method is less precise, and more variable compared to using oxygen uptake (VO_2_) reserve when prescribing exercise intensity. Indeed, each method has advantages and limitations (Mann et al., 2013). We decided to prescribe the exercise at a percentage of HRR for two reasons. Firstly, this approach accounts for individual differences and potential inaccuracies associated with heart rate-dependent methods or oxygen uptake methods such as percent of peak heart rate (%HR_peak_) or peak oxygen uptake (%VO_2peak_). Therefore, %HRR provides less risk of overestimating or underestimating exercise intensity (ACSM 2018). Secondly, %HRR allows for easy translation of exercise prescriptions into practical use, as individuals can monitor their heart rate during exercise without the need for complex equipment or measurements. Finally, TMS measures during the Pre, Post_0_, and Post_20_ time points took 10-20 minutes, which could be considered a long period of time. Since exercise-induced M1 excitability modulation can be time-sensitive, it is possible that some subtle changes were not captured. Prior research has shown that the timing of TMS assessments is crucial for detecting short-lived changes in M1 excitable output (Ridding & Ziemann, 2010). Future studies could consider reducing the number of TMS measures collected post-exercise to further understand the time-sensitive effects following acute aerobic exercise.

## Conclusion

This study investigated the impact of acute aerobic exercise intensity on distinct motor cortical circuits. Our main findings indicate that HIIT increased M1 excitable output to the greatest extent, and MIIT reduced GABAergic inhibition as measured by SICI to the greatest extent. Importantly, MIIT also transiently increased M1 excitable output, and LIIT, MIIT, and HIIT reduced GABAergic inhibition transiently. There is also evidence for preferential modulation of interneurons sensitive to anterior-to-posterior TMS current interacting with SICI, particularly following MIIT. Our findings provide evidence for a nuanced dose-response effect of exercise intensity on the modulation of distinct motor cortical circuits.

## Additional information

## Data availability statement

All main data presented in this manuscript are included within the figures. The datasets produced and analyzed in this study are available from the corresponding author upon reasonable request.

## Author contributions

NH contributed to the methodological development of the study, collected and analyzed the data and wrote the first draft of the manuscript. AO’F, LY, LB, HM, and MJ contributed to data collection. LB and MJ contributed to data processing. JLN and BP conceived of the project, contributed to interpretation of data as well as writing and editing the manuscript. All authors edited the manuscript and approved the final version before submission.

## Acknowledgements

We thank Callum O’Malley and Jonathan Tremblay for their valuable advice during the conception of this project. We also thank Maxime Bergevin for contributing to data collection and statistical analysis. Finally, we thank Yasmine Mahrez for her contribution to data processing.

## REFERENCES

ACSM. (2018). ACSM’s Guidelines for Exercise Testing and Prescription. Wolters Kluwer. https://books.google.ca/books?id=m_L-jwEACAAJ

Amassian, V., Stewart, M., Quirk, G., & Rosenthal, J. (1987). Physiological basis of motor effects of a transient stimulus to cerebral cortex. Neurosurgery, 20(1), 74–93.

Andrews, S. C., Curtin, D., Hawi, Z., Wongtrakun, J., Stout, J. C., & Coxon, J. P. (2020). Intensity Matters: High-intensity Interval Exercise Enhances Motor Cortex Plasticity More Than Moderate Exercise. Cereb Cortex, 30(1), 101–112. 10.1093/cercor/bhz075

Beneke, R., Hutler, M., Von Duvillard, S. P., Sellens, M., & Leithauser, R. M. (2003). Effect of test interruptions on blood lactate during constant workload testing. Med Sci Sports Exerc, 35(9), 1626–1630. 10.1249/01.Mss.0000084520.80451.D5

Booth, M. (2000). Assessment of Physical Activity: An International Perspective. Research Quarterly for Exercise and Sport, 71(sup2), 114–120. 10.1080/02701367.2000.11082794

Borg, E., & Kaijser, L. (2006). A comparison between three rating scales for perceived exertion and two different work tests. Scand J Med Sci Sports, 16(1), 57–69. 10.1111/j.1600-0838.2005.00448.x

Brown, K. E., Neva, J. L., Mang, C. S., Chau, B., Chiu, L. K., Francisco, B. A., Staines, W. R., & Boyd, L. A. (2020). The influence of an acute bout of moderate-intensity cycling exercise on sensorimotor integration. European Journal of Neuroscience, 52(12), 4779–4790.

Cirillo, J., & Byblow, W. D. (2016). Threshold tracking primary motor cortex inhibition: the influence of current direction. Eur J Neurosci, 44(8), 2614–2621. 10.1111/ejn.13369

Cirillo, J., Semmler, J. G., Mooney, R. A., & Byblow, W. D. (2018). Conventional or threshold-hunting TMS? A tale of two SICIs. Brain Stimulation, 11(6), 1296–1305. 10.1016/j.brs.2018.07.047

Coco, M., Alagona, G., Perciavalle, V., Perciavalle, V., Cavallari, P., & Caronni, A. (2014). Changes in cortical excitability and blood lactate after a fatiguing hand-grip exercise. Somatosens Mot Res, 31(1), 35–39. 10.3109/08990220.2013.834816

Coco, M., Alagona, G., Rapisarda, G., Costanzo, E., Calogero, R. A., Perciavalle, V., & Perciavalle, V. (2010). Elevated blood lactate is associated with increased motor cortex excitability. Somatosens Mot Res, 27(1), 1–8. 10.3109/08990220903471765

Coco, M., Buscemi, A., Ramaci, T., Tusak, M., Corrado, D. D., Perciavalle, V., Maugeri, G., Perciavalle, V., & Musumeci, G. (2020). Influences of Blood Lactate Levels on Cognitive Domains and Physical Health during a Sports Stress. Brief Review. Int J Environ Res Public Health, 17(23). 10.3390/ijerph17239043

Cohen, J. (2013). Statistical power analysis for the behavioral sciences. Routledge.

Craig, C. L., Marshall, A. L., Sjöström, M., Bauman, A. E., Booth, M. L., Ainsworth, B. E., Pratt, M., Ekelund, U., Yngve, A., Sallis, J. F., & Oja, P. (2003). International physical activity questionnaire: 12-country reliability and validity. Med Sci Sports Exerc, 35(8), 1381–1395. 10.1249/01.Mss.0000078924.61453.Fb

Curtin, D., Taylor, E. M., Bellgrove, M. A., Chong, T. T. J., & Coxon, J. P. (2023). D2 receptor blockade eliminates exercise-induced changes in cortical inhibition and excitation. Brain Stimulation, 16(3), 727–733. 10.1016/j.brs.2023.04.019

Day, B., Dressler, D., Maertens de Noordhout, A., Marsden, C., Nakashima, K., Rothwell, J., & Thompson, P. (1989). Electric and magnetic stimulation of human motor cortex: surface EMG and single motor unit responses. The Journal of physiology, 412(1), 449–473.

Di Lazzaro, V., Oliviero, A., Saturno, E., Pilato, F., Insola, A., Mazzone, P., Profice, P., Tonali, P., & Rothwell, J. C. (2001). The effect on corticospinal volleys of reversing the direction of current induced in the motor cortex by transcranial magnetic stimulation. Exp Brain Res, 138(2), 268–273. 10.1007/s002210100722

Di Lazzaro, V., Pilato, F., Dileone, M., Profice, P., Ranieri, F., Ricci, V., Bria, P., Tonali, P. A., & Ziemann, U. (2007). Segregating two inhibitory circuits in human motor cortex at the level of GABAA receptor subtypes: a TMS study. Clin Neurophysiol, 118(10), 2207–2214. 10.1016/j.clinph.2007.07.005

Di Lazzaro, V., Profice, P., Ranieri, F., Capone, F., Dileone, M., Oliviero, A., & Pilato, F. (2012). I-wave origin and modulation. Brain Stimul, 5(4), 512–525. 10.1016/j.brs.2011.07.008

El-Sayes, J., Turco, C. V., Skelly, L. E., Locke, M. B., Gibala, M. J., & Nelson, A. J. (2020). Acute high-intensity and moderate-intensity interval exercise do not change corticospinal excitability in low fit, young adults. PLoS ONE, 15 (*1*) (no pagination)(e0227581). 10.1371/journal.pone.0227581

El-Sayes, J., Turco, C. V., Skelly, L. E., Nicolini, C., Fahnestock, M., Gibala, M. J., & Nelson, A. J. (2019). The Effects of Biological Sex and Ovarian Hormones on Exercise-Induced Neuroplasticity. Neuroscience, 410, 29–40. 10.1016/j.neuroscience.2019.04.054

Ferrer-Uris, B., Busquets, A., Lopez-Alonso, V., Fernandez-Del-Olmo, M., & Angulo-Barroso, R. (2017). Enhancing consolidation of a rotational visuomotor adaptation task through acute exercise. PLoS One, 12(4), e0175296. 10.1371/journal.pone.0175296

Flores-Hernandez, J., Hernandez, S., Snyder, G. L., Yan, Z., Fienberg, A. A., Moss, S. J., Greengard, P., & Surmeier, D. J. (2000). D(1) dopamine receptor activation reduces GABA(A) receptor currents in neostriatal neurons through a PKA/DARPP-32/PP1 signaling cascade. J Neurophysiol, 83(5), 2996–3004. 10.1152/jn.2000.83.5.2996

Garnier, Y. M., Lepers, R., Dubau, Q., Pageaux, B., & Paizis, C. (2018). Neuromuscular and perceptual responses to moderate-intensity incline, level and decline treadmill exercise. European Journal of Applied Physiology, 118, 2039–2053.

Greenwood, B. N. (2019). The role of dopamine in overcoming aversion with exercise. Brain Res, 1713, 102–108. 10.1016/j.brainres.2018.08.030

Grossl, T., de Lucas, R. D., de Souza, K. M., & Guglielmo, L. G. A. (2012). Time to exhaustion at intermittent maximal lactate steady state is longer than continuous cycling exercise. Applied Physiology, Nutrition, and Metabolism, 37(6), 1047–1053.

Hamada, M., Galea, J. M., Di Lazzaro, V., Mazzone, P., Ziemann, U., & Rothwell, J. C. (2014). Two distinct interneuron circuits in human motor cortex are linked to different subsets of physiological and behavioral plasticity. J Neurosci, 34(38), 12837–12849. 10.1523/jneurosci.1960-14.2014

Hamada, M., Murase, N., Hasan, A., Balaratnam, M., & Rothwell, J. C. (2013). The role of interneuron networks in driving human motor cortical plasticity. Cereb Cortex, 23(7), 1593–1605. 10.1093/cercor/bhs147

Hanajima, R., Ugawa, Y., Terao, Y., Sakai, K., Furubayashi, T., Machii, K., & Kanazawa, I. (1998). Paired-pulse magnetic stimulation of the human motor cortex: differences among I waves. J Physiol, 509 (*Pt 2*)(Pt 2), 607–618. 10.1111/j.1469-7793.1998.607bn.x

Hannah, R., Cavanagh, S. E., Tremblay, S., Simeoni, S., & Rothwell, J. C. (2018). Selective Suppression of Local Interneuron Circuits in Human Motor Cortex Contributes to Movement Preparation. J Neurosci, 38(5), 1264–1276. 10.1523/jneurosci.2869-17.2017

Hardy, C. J., & Rejeski, W. J. (1989). Not what, but how one feels: the measurement of affect during exercise. Journal of sport and exercise psychology, 11(3), 304–317.

Hendy, A. M., Andrushko, J. W., Della Gatta, P. A., & Teo, W.-P. (2022). Acute effects of high-intensity aerobic exercise on motor cortical excitability and inhibition in sedentary adults. Frontiers in Psychology, 13, 1192.

Kaneko, K., Kawai, S., Fuchigami, Y., Morita, H., & Ofuji, A. (1996). The effect of current direction induced by transcranial magnetic stimulation on the corticospinal excitability in human brain. Electroencephalography and clinical neurophysiology/electromyography and motor control, 101(6), 478–482.

Kann, O. (2024). Lactate as a supplemental fuel for synaptic transmission and neuronal network oscillations: Potentials and limitations. J Neurochem, 168(5), 608–631. 10.1111/jnc.15867

Kanthack, T. F. D., Guillot, A., Clémençon, M., Debarnot, U., & Di Rienzo, F. (2020). Effect of physical fatigue elicited by continuous and intermittent exercise on motor imagery ability. Research Quarterly for Exercise and Sport, 91(3), 525–538.

Karvonen, J., & Vuorimaa, T. (1988). Heart rate and exercise intensity during sports activities. Practical application. Sports Medicine, 5(5), 303–311. 10.2165/00007256-198805050-00002

Kuo, Hsieh, M. H., Lin, Y. T., Kuo, M. F., & Nitsche, M. A. (2023). A single bout of aerobic exercise modulates motor learning performance and cortical excitability in humans. Int J Clin Health Psychol, 23(1), 100333. 10.1016/j.ijchp.2022.100333

Lakens, D. (2022). Sample size justification. Collabra: psychology, 8(1), 33267.

Lazzaro, V. D., Restuccia, D., Oliviero, A., Profice, P., Ferrara, L., Insola, A., Mazzone, P., Tonali, P., & Rothwell, J. (1998). Magnetic transcranial stimulation at intensities below active motor threshold activates intracortical inhibitory circuits. Experimental brain research, 119, 265–268.

Lulic, T., El-Sayes, J., Fassett, H. J., & Nelson, A. J. (2017). Physical activity levels determine exercise-induced changes in brain excitability. PLoS One, 12(3), e0173672. 10.1371/journal.pone.0173672

MacDonald, M. A., Khan, H., Kraeutner, S. N., Usai, F., Rogers, E. A., Kimmerly, D. S., Dechman, G., & Boe, S. G. (2019). Intensity of acute aerobic exercise but not aerobic fitness impacts on corticospinal excitability [Clinical Trial]. *Applied Physiology, Nutrition, & Metabolism = Physiologie Appliquee*, Nutrition et Metabolisme, 44(8), 869–878. 10.1139/apnm-2018-0643

Mang, C. S., Campbell, K. L., Ross, C. J., & Boyd, L. A. (2013). Promoting neuroplasticity for motor rehabilitation after stroke: considering the effects of aerobic exercise and genetic variation on brain-derived neurotrophic factor. Phys Ther, 93(12), 1707–1716. 10.2522/ptj.20130053

Mang, C. S., Snow, N. J., Campbell, K. L., Ross, C. J., & Boyd, L. A. (2014). A single bout of high-intensity aerobic exercise facilitates response to paired associative stimulation and promotes sequence-specific implicit motor learning [Research Support, Non-U.S. Gov’t]. Journal of Applied Physiology, 117(11), 1325–1336. 10.1152/japplphysiol.00498.2014

Mann, T., Lamberts, R. P., & Lambert, M. I. (2013). Methods of prescribing relative exercise intensity: physiological and practical considerations. Sports Med, 43(7), 613–625. 10.1007/s40279-013-0045-x

McDonnell, M. N., Buckley, J. D., Opie, G. M., Ridding, M. C., & Semmler, J. G. (2013). A single bout of aerobic exercise promotes motor cortical neuroplasticity [Research Support, Non-U.S. Gov’t]. Journal of Applied Physiology, 114(9), 1174–1182. 10.1152/japplphysiol.01378.2012

Merrell, L. H., Perkin, O. J., Bradshaw, L., Collier-Bain, H. D., Collins, A. J., Davies, S., Eddy, R., Hickman, J. A., Nicholas, A. P., Rees, D., Spellanzon, B., James, L. J., McKay, A. K. A., Smith, H. A., Turner, J. E., Koumanov, F., Maher, J., Thompson, D., Gonzalez, J. T., & Betts, J. A. (2024). Myths and Methodologies: Standardisation in human physiology research-should we control the controllables? Exp Physiol, 109(7), 1099–1108. 10.1113/ep091557

Mirdamadi, J. L., Suzuki, L. Y., & Meehan, S. K. (2017). Attention modulates specific motor cortical circuits recruited by transcranial magnetic stimulation. Neuroscience, 359, 151–158. 10.1016/j.neuroscience.2017.07.028

Mooney, R. A., Coxon, J. P., Cirillo, J., Glenny, H., Gant, N., & Byblow, W. D. (2016). Acute aerobic exercise modulates primary motor cortex inhibition. Exp Brain Res, 234(12), 3669–3676. 10.1007/s00221-016-4767-5

Morris, T. P., Fried, P. J., Macone, J., Stillman, A., Gomes-Osman, J., Costa-Miserachs, D., Tormos Muñoz, J. M., Santarnecchi, E., & Pascual-Leone, A. (2020). Light aerobic exercise modulates executive function and cortical excitability. Eur J Neurosci, 51(7), 1723–1734. 10.1111/ejn.14593

Myers, J., Arena, R., Franklin, B., Pina, I., Kraus, W. E., McInnis, K., & Balady, G. J. (2009). Recommendations for clinical exercise laboratories: a scientific statement from the American Heart Association. Circulation, 119(24), 3144–3161.

Neva, J. L., Brown, K. E., Mang, C. S., Francisco, B. A., & Boyd, L. A. (2017). An acute bout of exercise modulates both intracortical and interhemispheric excitability. European Journal of Neuroscience, 45(10), 1343–1355. 10.1111/ejn.13569

Neva, J. L., Brown, K. E., Peters, S., Feldman, S. J., Mahendran, N., Boisgontier, M. P., & Boyd, L. A. (2021). Acute Exercise Modulates the Excitability of Specific Interneurons in Human Motor Cortex. Neuroscience, 475, 103–116. 10.1016/j.neuroscience.2021.08.032

Ni, Z., Charab, S., Gunraj, C., Nelson, A. J., Udupa, K., Yeh, I.-J., & Chen, R. (2011). Transcranial magnetic stimulation in different current directions activates separate cortical circuits. Journal of Neurophysiology, 105(2), 749–756.

Nicolini, C., Michalski, B., Toepp, S. L., Turco, C. V., D’Hoine, T., Harasym, D., Gibala, M. J., Fahnestock, M., & Nelson, A. J. (2020). A Single Bout of High-intensity Interval Exercise Increases Corticospinal Excitability, Brain-derived Neurotrophic Factor, and Uncarboxylated Osteolcalcin in Sedentary, Healthy Males. Neuroscience, 437, 242–255. 10.1016/j.neuroscience.2020.03.042

Oldfield, R. C. (1971). The assessment and analysis of handedness: the Edinburgh inventory. Neuropsychologia, 9(1), 97–113. 10.1016/0028-3932(71)90067-4

Opie, G. M., & Semmler, J. G. (2019). Acute Exercise at Different Intensities Influences Corticomotor Excitability and Performance of a Ballistic Thumb Training Task. Neuroscience, 412, 29–39. 10.1016/j.neuroscience.2019.05.049

Ostadan, F., Centeno, C., Daloze, J. F., Frenn, M., Lundbye-Jensen, J., & Roig, M. (2016). Changes in corticospinal excitability during consolidation predict acute exercise-induced off-line gains in procedural memory. Neurobiology of Learning & Memory, 136, 196–203. 10.1016/j.nlm.2016.10.009

Paulus, W., Classen, J., Cohen, L. G., Large, C. H., Di Lazzaro, V., Nitsche, M., Pascual-Leone, A., Rosenow, F., Rothwell, J. C., & Ziemann, U. (2008). State of the art: Pharmacologic effects on cortical excitability measures tested by transcranial magnetic stimulation. Brain Stimul, 1(3), 151–163. 10.1016/j.brs.2008.06.002

Peurala, S. H., Müller-Dahlhaus, J. F., Arai, N., & Ziemann, U. (2008). Interference of short-interval intracortical inhibition (SICI) and short-interval intracortical facilitation (SICF). Clin Neurophysiol, 119(10), 2291–2297. 10.1016/j.clinph.2008.05.031

Piercy, K. L., Troiano, R. P., Ballard, R. M., Carlson, S. A., Fulton, J. E., Galuska, D. A., George, S. M., & Olson, R. D. (2018). The Physical Activity Guidelines for Americans. Jama, 320(19), 2020–2028. 10.1001/jama.2018.14854

Roig, M., Skriver, K., Lundbye-Jensen, J., Kiens, B., & Nielsen, J. B. (2012). A single bout of exercise improves motor memory.

Sacheli, M. A., Neva, J. L., Lakhani, B., Murray, D. K., Vafai, N., Shahinfard, E., English, C., McCormick, S., Dinelle, K., Neilson, N., McKenzie, J., Schulzer, M., McKenzie, D. C., Appel-Cresswell, S., McKeown, M. J., Boyd, L. A., Sossi, V., & Stoessl, A. J. (2019). Exercise increases caudate dopamine release and ventral striatal activation in Parkinson’s disease. Mov Disord, 34(12), 1891–1900. 10.1002/mds.27865

Safikhani, S., Gries, K. S., Trudeau, J. J., Reasner, D., Rüdell, K., Coons, S. J., Bush, E. N., Hanlon, J., Abraham, L., & Vernon, M. (2018). Response scale selection in adult pain measures: results from a literature review. Journal of patient-reported outcomes, 2, 1–9.

Sakai, K., Ugawa, Y., Terao, Y., Hanajima, R., Furubayashi, T., & Kanazawa, I. (1997). Preferential activation of different I waves by transcranial magnetic stimulation with a figure-of-eight-shaped coil. Experimental Brain Research, 113(1), 24–32. 10.1007/BF02454139

Sale, Lavender, A. P., Opie, G. M., Nordstrom, M. A., & Semmler, J. G. (2016). Increased intracortical inhibition in elderly adults with anterior–posterior current flow: A TMS study. Clinical Neurophysiology, 127(1), 635–640.

Sale, M. V., Ridding, M. C., & Nordstrom, M. A. (2008). Cortisol inhibits neuroplasticity induction in human motor cortex. J Neurosci, 28(33), 8285–8293. 10.1523/jneurosci.1963-08.2008

Singh, Duncan, R. E., Neva, J. L., & Staines, W. R. (2014). Aerobic exercise modulates intracortical inhibition and facilitation in a nonexercised upper limb muscle. BMC sports science, medicine and rehabilitation, 6, 1–10.

Singh, A. M., Neva, J. L., & Staines, W. R. (2016). Aerobic exercise enhances neural correlates of motor skill learning. Behav Brain Res, 301, 19–26. 10.1016/j.bbr.2015.12.020

Skriver, K., Roig, M., Lundbye-Jensen, J., Pingel, J., Helge, J. W., Kiens, B., & Nielsen, J. B. (2014). Acute exercise improves motor memory: exploring potential biomarkers. Neurobiol Learn Mem, 116, 46–58. 10.1016/j.nlm.2014.08.004

Smith, A. E., Goldsworthy, M. R., Garside, T., Wood, F. M., & Ridding, M. C. (2014). The influence of a single bout of aerobic exercise on short-interval intracortical excitability. Exp Brain Res, 232(6), 1875–1882. 10.1007/s00221-014-3879-z

Smith, A. E., Goldsworthy, M. R., Wood, F. M., Olds, T. S., Garside, T., & Ridding, M. C. (2018). High-intensity Aerobic Exercise Blocks the Facilitation of iTBS-induced Plasticity in the Human Motor Cortex. Neuroscience, 373, 1–6. 10.1016/j.neuroscience.2017.12.034

Spampinato, D. (2020). Dissecting two distinct interneuronal networks in M1 with transcranial magnetic stimulation. Exp Brain Res, 238(7-8), 1693–1700. 10.1007/s00221-020-05875-y

Spampinato, D., Ibáñez, J., Spanoudakis, M., Hammond, P., & Rothwell, J. C. (2020). Cerebellar transcranial magnetic stimulation: The role of coil type from distinct manufacturers. Brain Stimulation, 13(1), 153–156. 10.1016/j.brs.2019.09.005

Stavrinos, E. L., & Coxon, J. P. (2017). High-intensity interval exercise promotes motor cortex disinhibition and early motor skill consolidation. Journal of Cognitive Neuroscience, 29(4), 593–604. 10.1162/jocn_a_01078

Tamm, A. S., Lagerquist, O., Ley, A. L., & Collins, D. F. (2009). Chronotype influences diurnal variations in the excitability of the human motor cortex and the ability to generate torque during a maximum voluntary contraction. Journal of Biological Rhythms, 24(3), 211–224.

Thomas, R., Johnsen, L. K., Geertsen, S. S., Christiansen, L., Ritz, C., Roig, M., & Lundbye-Jensen, J. (2016). Acute Exercise and Motor Memory Consolidation: The Role of Exercise Intensity. PLoS One, 11(7), e0159589. 10.1371/journal.pone.0159589

Tschakert, G., & Hofmann, P. (2013). High-intensity intermittent exercise: methodological and physiological aspects. Int J Sports Physiol Perform, 8(6), 600–610. 10.1123/ijspp.8.6.600

Turco, C. V., & Nelson, A. J. (2021). Transcranial Magnetic Stimulation to Assess Exercise-Induced Neuroplasticity. Frontiers in Neuroergonomics,

Yamazaki, Y., Sato, D., Yamashiro, K., Nakano, S., Onishi, H., & Maruyama, A. (2019). Acute Low-Intensity Aerobic Exercise Modulates Intracortical Inhibitory and Excitatory Circuits in an Exercised and a Non-exercised Muscle in the Primary Motor Cortex. Frontiers in Physiology, 10 (no pagination)(1361). 10.3389/fphys.2019.01361

Youssef, L., Harroum, N., Francisco, B. A., Johnson, L., Arvisais, D., Pageaux, B., Romain, A. J., Hayward, K. S., & Neva, J. L. (2024). Neurophysiological effects of acute aerobic exercise in young adults: a systematic review and meta-analysis. Neurosci Biobehav Rev, 164, 105811. 10.1016/j.neubiorev.2024.105811

Ziemann, U., & Rothwell, J. C. (2000). I-waves in motor cortex. Journal of Clinical Neurophysiology, 17(4), 397–405.

Zuniga, J. M., Berg, K., Noble, J., Harder, J., Chaffin, M. E., & Hanumanthu, V. S. (2011). Physiological Responses during Interval Training with Different Intensities and Duration of Exercise. The Journal of Strength & Conditioning Research, 25(5), 1279–1284. 10.1519/JSC.0b013e3181d681b6

