## Supporting_information_Fig1 for "Motor cortical circuits are uniquely impacted by different exercise intensities"

**Supporting Information – Fig. 1: Changes in heart rate and perceptual responses to the three intermittent cycling exercise and the rest condition.**

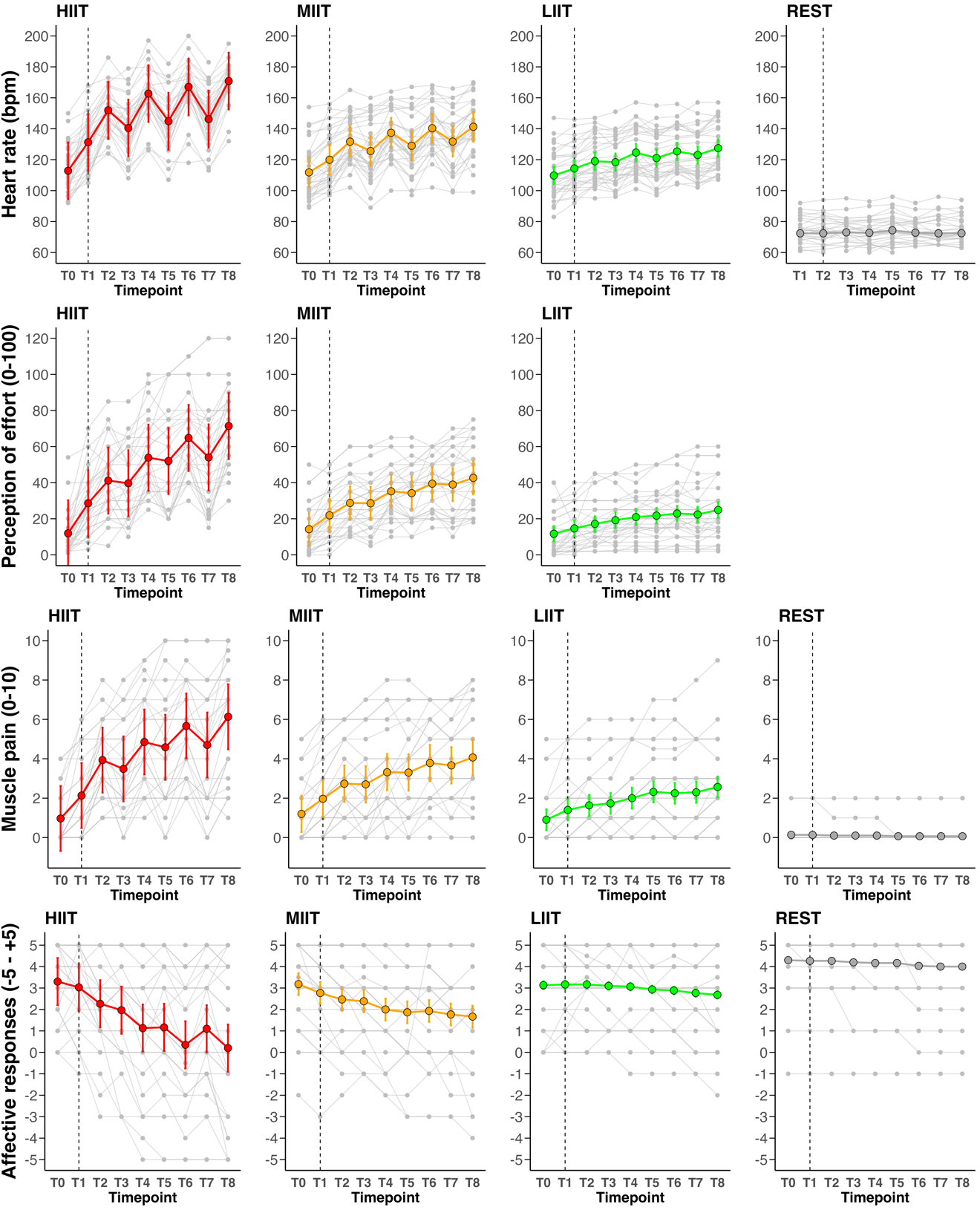

***Caption for* Supporting Information – *Fig. 1****:*

*The large circles represent the average for each condition with dark grey for Rest, green for LIIT, orange for MIIT and red for HIIT, the light grey dots represent the individual data.* ***(A)*** *Displays the raw heart rate data.* ***(B)*** *Displays the perception of effort data.* ***(C)*** *Displays the muscle pain data. (****D)*** *Displays the affective responses data.*

*Abbreviations:* ***bpm****: beats per minute;* ***HIIT****: high intensity interval training session;* ***LIIT****: light intensity interval training session;* ***MIIT****: moderate intensity interval training session;* ***T0****: during the last 20 seconds of the 5-minute-warm up,* ***T1****: during the first 20 seconds of the block 1,* ***T2****: at 2 minutes 30 seconds of block 1;* ***T3****: during the first 20 seconds of the block 2;* ***T4****: at 2 minutes 30 seconds of block 2;* ***T5****: during the first 20 seconds of the block 3;* ***T6****: at 2 minutes 30 seconds of block 3;* ***T7****: during the first 20 seconds of the block 4;* ***T8****: at 2 minutes 30 seconds of block 4.*
