## Supporting_information_Table1 for "Motor cortical circuits are uniquely impacted by different exercise intensities"

**Supporting Information – *Table 1: Difference in cycling power output, heart rate and perceptual responses between the warm-up and the intervention in all four conditions.***

|  | ***HIIT*** | ***MIIT*** | ***LIIT*** | ***REST*** |
| --- | --- | --- | --- | --- |
| ***Power output (% peak value)*** |  |  |  |  |
| Warm-up/ active recovery | *27 (9)* | *27 (9)* | *27 (9)* | *NA* |
| Intervention | *83 (12)* | *58 (9)* | *38 (8)* | *NA* |
| ***Heart Rate (% peak value)*** |  |  |  |  |
| Warm-up | *63 (9)* | *63 (13)* | *61 (10)* | *41 (7)* |
| Intervention | *85 (8)* | *74 (9)* | *68 (10)* | *41 (6)* |
| ***Perception of effort (0-100)*** |  |  |  |  |
| Warm-up | *12 (12)* | *18 (20)* | *12 (12)* | *NA* |
| Intervention | *51 (14)* | *34 (6)* | *20 (3)* | *NA* |
| ***Muscle pain (0-10)*** |  |  |  |  |
| Warm-up | *1 (1)* | *4 (1)* | *3 (2)* | *0 (1)* |
| Intervention | *4 (1)* | *3 (1)* | *2 (0.4)* | *0 (0.02)* |
| ***Affective responses (-5 - +5)*** |  |  |  |  |
| Warm-up | *3 (2)* | *3 (2)* | *3 (2)* | *4 (1)* |
| Intervention | *1 (1)* | *2 (0.4)* | *3 (0.2)* | *4 (0.1)* |

***Caption for* Supporting Information – *Table 1:***

*All data are reported as mean (SD).* *Abbreviations:* ***Active recovery:*** *during the 2 minutes active following the 3-minute interval of cycling at the target intensity;* ***HIIT****: high intensity interval training session; I****ntervention****: average data during the 20 min of cycling exercise or rest;* ***LIIT****: light intensity interval training session;* ***MIIT****: moderate intensity interval training session;* ***NA****: not assessed;* ***Warm-up****: during the last 20 seconds of the 5-minute-warm up or rest.*
