## Supporting_information_Table2 for "Motor cortical circuits are uniquely impacted by different exercise intensities"

**Supporting Information – *Table 2: Detailed statistical results for heart rate, perception of effort, muscle pain and affective responses***

|  | | | ***df*** | | ***F*** | ***p.value*** | | | ***η²_p_*** |
| --- | --- | --- | --- | --- | --- | --- | --- | --- | --- |
| ***Heart rate (% peak value)*** | Condition | 3, 87 | | 287.3 | | |  | **< .001** | 0.91 |
|  | Time | 3, 87 | | 64.14 | | |  | **< .001** | 0.69 |
|  | Condition * Time | 9, 261 | | 14.45 | | |  | **< .001** | 0.30 |
| ***Perception of effort*** | Condition | 2, 58 | | 51.56 | | |  | **< .001** | 0.64 |
|  | Time | 3, 87 | | 60.72 | | |  | **< .001** | 0.68 |
|  | Condition * Time | 6, 174 | | 16.47 | | |  | **< .001** | 0.36 |
| ***Muscle pain*** | Condition | 3, 87 | | 59.45 | | |  | **< .001** | 0.67 |
|  | Time | 3, 87 | | 34.45 | | |  | **< .001** | 0.54 |
|  | Condition * Time | 9, 261 | | 12.27 | | |  | **< .001** | 0.30 |
| ***Affective responses*** | Condition | 3, 87 | | 23.23 | | |  | **< .001** | 0.44 |
|  | Time | 3, 87 | | 19.03 | | |  | **< .001** | 0.40 |
|  | Condition * Time | 9, 261 | | 6.69 | | |  | **< .001** | 0.19 |

***Caption for* Supporting Information – *Table 2:***

*Abbreviations:* ***HIIT****: high intensity interval training session;* ***MIIT****: moderate intensity interval training session;* ***LIIT****: light intensity interval training session;* ***df****: degree of freedom;* ***η²_p_****: eta square partial.*
