## Supporting_information_Table3 for "Motor cortical circuits are uniquely impacted by different exercise intensities"

**Supporting Information – *Table* *3: Post-hocs for exercise-related data***

|  | |  | ***Heart rate (% peak value)*** | ***Perception of effort*** | ***Muscle pain*** | ***Affective responses*** |
| --- | --- | --- | --- | --- | --- | --- |
| ***Comparison*** | | ***Time*** | ***p.value*** | | | |
| HIIT | - MIIT | B1 | **< .001** | **.017** | **.030** | .119 |
| HIIT | - LIIT | B1 | **< .001** | **< .001** | **< .001** | .702 |
| HIIT | - REST | B1 | **< .001** | **-** | **< .001** | .132 |
| MIIT | - LIIT | B1 | **.003** | **< .001** | **.006** | .073 |
| MIIT | - REST | B1 | **< .001** | **-** | **< .001** | **< .001** |
| LIIT | - REST | B1 | **< .001** | **-** | **< .001** | **.009** |
| HIIT | - MIIT | B2 | **.003** | **.002** | **.017** | .140 |
| HIIT | - LIIT | B2 | **< .001** | **< .001** | **< .001** | **.004** |
| HIIT | - REST | B2 | **< .001** | **-** | **< .001** | **< .001** |
| MIIT | - LIIT | B2 | **< .001** | **< .001** | **.023** | **.043** |
| MIIT | - REST | B2 | **< .001** | **-** | **< .001** | **< .001** |
| LIIT | - REST | B2 | **< .001** | **-** | **< .001** | **.023** |
| HIIT | - MIIT | B3 | **< .001** | **< .001** | **.008** | .119 |
| HIIT | - LIIT | B3 | **< .001** | **< .001** | **< .001** | **< .001** |
| HIIT | - REST | B3 | **< .001** | **-** | **< .001** | **< .001** |
| MIIT | - LIIT | B3 | **< .001** | **< .001** | **.01** | **.006** |
| MIIT | - REST | B3 | **< .001** | **-** | **< .001** | **< .001** |
| LIIT | - REST | B3 | **< .001** | **-** | **< .001** | **.023** |
| HIIT | - MIIT | B4 | **< .001** | **< .001** | **.019** | .151 |
| HIIT | - LIIT | B4 | **< .001** | **< .001** | **< .001** | **.004** |
| HIIT | - REST | B4 | **< .001** | **-** | **< .001** | **< .001** |
| MIIT | - LIIT | B4 | **< .001** | **< .001** | **.007** | **.022** |
| MIIT | - REST | B4 | **< .001** | **-** | **< .001** | **.002** |
| LIIT | - REST | B4 | **< .001** | **-** | **< .001** | **.041** |

***Caption for* Supporting Information – *Table 3:***

*Post hoc results are reported with Holm-Bonferroni correction. Abbreviations:* ***HIIT****: high intensity interval training session;* ***MIIT****: moderate intensity interval training session;* ***LIIT****: light intensity interval training session;* ***B1****: block 1 (0-3 minutes);* ***B2****: block 2 (5-8 minutes);* ***B3****: block 3 (10-13 minutes);* ***B4****: block 4 (15-18 minutes).*
