## Supporting_information_Table6 for "Motor cortical circuits are uniquely impacted by different exercise intensities"

**Supporting Information – *Table 6: Test Stimulus motor evoked potential (MEP) data***

| *TMS cuurent* | *Condition* | ***Pre*** | ***Post_0_*** | ***Post_20_*** |
| --- | --- | --- | --- | --- |
| *Posterior-to-anterior current (PA)* | HIIT | 1.13 (0.16) | 1.15 (0.19) | 1.1 (0.19) |
|  | MIIT | 1.14 (0.21) | 1.13 (0.2) | 1.13 (0.2) |
|  | LIIT | 1.14 (0.16) | 1.12 (0.17) | 1.13 (0.15) |
|  | REST | 1.23 (0.21) | 1.29 (0.19) | 1.26 (0.2) |
| *Anterior-to-posterior current (AP)* | HIIT | 1.14 (0.16) | 1.15 (0.19) | 1.14 (0.24) |
|  | MIIT | 1.15 (0.21) | 1.16 (0.22) | 1.13 (0.2) |
|  | LIIT | 1.14 (0.16) | 1.13 (0.16) | 1.13 (0.14) |
|  | REST | 1.24 (0.2) | 1.27 (0.18) | 1.27 (0.2) |

***Caption for* Supporting Information – *Table 6:***

*Data shown is the peak-to-peak amplitude of the motor evoked potential expressed in mV and reported as mean (SD).* *Abbreviations:* ***HIIT****: high intensity interval training session;* ***LIIT****: light intensity interval training session;* ***MIIT****: moderate intensity interval training session;* ***Post_0_****: immediately after exercise/rest;* ***Post_20_****: 20 minutes after exercise/rest;* ***Pre****: before exercise/rest;* ***TMS****: transcranial magnetic stimulation.*
