## Supporting_information_Table7 for "Motor cortical circuits are uniquely impacted by different exercise intensities"

**Supporting Information – *Table 7: SICI results for pairwise comparisons for MIIT versus REST in PA and AP TMS currents***

|  | ***Condition*** | ***Timepoint*** | ***Condition*** | ***Timepoint*** | ***statistic*** | ***p.value*** | ***Effect size (Cohen d)*** |
| --- | --- | --- | --- | --- | --- | --- | --- |
| *Posterior-to-anterior current (PA)* | MIIT | ***Pre*** | REST | ***Pre*** | -3.12 | .057 | -0.57 |
|  |  |  | MIIT | ***Post_0_*** | -5.12 | **< .001** | -0.93 |
|  |  |  | MIIT | ***Post_20_*** | -4.95 | **< .001** | -0.9 |
|  |  | ***Post_0_*** | REST | ***Post_0_*** | 2.2 | .491 | 0.4 |
|  |  | ***Post_20_*** | REST | ***Post_20_*** | 1.39 | .208 | 0.23 |
|  | REST | ***Pre*** | REST | ***Post_0_*** | -1.59 | .121 | -0.29 |
|  |  |  | REST | ***Post_20_*** | -1.29 | .205 | -0.24 |
| *Anterior-to-posterior current (AP)* | MIIT | ***Pre*** | REST | ***Pre*** | -0.2 | .842 | -0.04 |
|  |  |  | MIIT | ***Post_0_*** | -4.2 | **.003** | -0.77 |
|  |  |  | MIIT | ***Post_20_*** | -4.36 | **.002** | -0.8 |
|  |  | ***Post_0_*** | REST | ***Post_0_*** | 3.45 | **.024** | 0.63 |
|  |  | ***Post_20_*** | REST | ***Post_20_*** | 3.59 | **.017** | 0.66 |
|  | REST | ***Pre*** | REST | ***Post_0_*** | -1.05 | .301 | -0.19 |
|  |  |  | REST | ***Post_20_*** | -0.9 | .375 | -0.16 |

***Caption for Supporting Information* – *Table 7:***

*Post hoc results are reported with Holm-Bonferroni correction. Abbreviations:* ***HIIT****: high intensity interval training session;* ***LIIT****: light intensity interval training session;* ***MIIT****: moderate intensity interval training session;* ***Post_0_****: immediately after exercise/rest;* ***Post_20_****: 20 minutes after exercise/rest;* ***Pre****: before exercise/rest;* ***TMS****: transcranial magnetic stimulation.*
