## Supporting_information_Table4 for "Motor cortical circuits are uniquely impacted by different exercise intensities"

**Supporting Information – *Table 4: Baseline resting motor threshold (RMT) data expressed as percentage of maximal stimulator output (%MSO)***

|  | Condition | Lateral-to-medial current (LM) | Posterior-to-anterior current (PA) | Anterior-to-posterior current (AP) |
| --- | --- | --- | --- | --- |
| *Pre-exercise/rest* | HIIT | 51 (6) | 46 (6) | 58 (9) |
|  | MIIT | 51 (6) | 45 (6) | 57 (9) |
|  | LIIT | 51 (6) | 45 (7) | 57 (10) |
|  | REST | 51 (7) | 45 (7) | 57 (9) |
| ***Overall Mean* ± SD** | | **51 (6)** | **45 (6)** | **57 (9)** |

***Caption for* Supporting Information – *Table 4:***

*Data shown is the percentage of maximal stimulator output (% MSO) reported as mean (SD).* *Abbreviations:* ***PA*** *posterior-to-anterior;* ***AP*** *anterior-to-posterior;* ***LM****: lateral-to-medial;* ***HIIT****: high intensity interval training session;* ***MIIT****: moderate intensity interval training session;* ***LIIT****: light intensity interval training session.*
